# Computational modeling of VEGF-A signaling predicts lymphatic junction state

**DOI:** 10.64898/2026.10.14.759242

**Authors:** Yiping Jia, Silvia Gonzalez-Nieves, Simon Guignard, Vincenza Cifarelli, Stacey D. Finley

## Abstract

Lymphatic endothelial cell (LEC) junctions switch between continuous zippers and discontinuous buttons to control fluid, cell and lipid uptake, but how the strength and timing of signaling set junction state is unknown. Here we present, to our knowledge, the first mechanistic signaling model calibrated to LEC data, linking VEGF-A-induced activation of VEGFR2 homodimers and VEGFR2/VEGFR3 heterodimers to the balance of active RAC1 and RhoA, summarized as a predicted zippering propensity. Calibrated to human dermal LEC time courses and validated against independent receptor knockdowns, the model predicts that VEGF-A dose primarily determines how long zippering-favoring signaling lasts following stimulation, and it identifies which of the parameters related to RAC1-RhoA kinetics control this duration. Using single-cell transcriptomes from high-fat-diet-fed mice, the model predicts that obesity shortens this response in mesenteric capillary and epididymal collector LECs but prolongs it in epididymal capillary LECs. Tuning RAC1–RhoA kinetics restores baseline in each population but requires opposite adjustments, suggesting that targeting this axis must be context-specific.

## INTRODUCTION

The lymphatic vasculature maintains tissue fluid balance, transports immune cells and absorbs dietary lipids. These functions depend on different permeability properties along the lymphatic tree; initial lymphatic capillaries take up fluid, cells and macromolecules, whereas collecting vessels retain lymph during transport^1^. These functional differences are supported in part by the organization of junctions between lymphatic endothelial cells (LECs). Lymphatic capillaries typically have discontinuous, button-like junctions that permit entry into the lumen, whereas collectors have continuous, zipper-like junctions that restrict paracellular passage^2,3^. Importantly, these junctions are dynamic and remodel during development and in response to inflammation, diet, and other local signals, allowing lymphatic permeability to adapt to changing physiological demands. Changes in junctional organization can therefore alter lymphatic function as excessive zippering can restrict uptake, whereas loss of junctional integrity can compromise lymph retention^4^.

VEGF-A–VEGFR signaling is an important regulator of lymphatic junctional plasticity. VEGF-A activates VEGFR2 homodimers and VEGFR2/VEGFR3 heterodimers, engaging downstream PI3K–AKT and small GTPase signaling. Within this network, RAC1 and RhoA exert opposing effects on endothelial junctions. RAC1 stabilizes VE-cadherin adhesions and limits junctional tension, whereas RhoA–ROCK signaling promotes actomyosin contractility and junctional remodeling^5–7^. The functional importance of this pathway has been demonstrated in intestinal lacteals, where increased VEGF-A–VEGFR2 activity promotes junction zippering and restricts chylomicron uptake, whereas attenuation of this pathway favors junction opening and lipid absorption^8–10^. However, most experimental studies assess signaling and junction organization at selected time points, limiting our understanding of how signaling evolves over time. An important unresolved question is how increasing VEGF-A stimulation changes the dynamics of junction-associated signaling.

Mechanistic computational models provide a framework for integrating these interactions and determine how receptor activation and downstream signaling kinetics shape the temporal response of LECs to VEGF-A. In blood endothelial cells, such models have been used to investigate VEGF–receptor interactions, receptor dimerization, trafficking, and downstream signaling^11–14^. More recently, these approaches have been extended to connect growth-factor signaling with endothelial permeability^15^. In contrast, computational studies of the lymphatic vasculature have focused primarily on lymph transport, vessel mechanics, valve function, and pumping^16^. A mechanistic model that links VEGF-A signaling dynamics to predicted junction behavior in LECs would provide a means to examine how receptor activation and downstream signaling collectively shape the junction-associated response over time.

Cell-to-cell heterogeneity adds another layer of complexity to signaling dynamics. Even within the same endothelial population, cells differ in the abundance of receptors and signaling proteins, which can influence the magnitude and duration of their responses to a common stimulus^17^. Models based on a single representative cell may therefore overlook differences in signaling behavior across a population. Incorporating variability in signaling components can help determine whether changes in VEGF-A stimulation produce uniform responses or distinct patterns of signaling activity among individual LECs. Beyond cell-to-cell variability, LECs exhibit molecular differences across tissues and vessel types that may influence their responses to VEGF-A. These differences are particularly relevant in obesity, which is associated with lymphatic dysfunction and remodeling of the lymphatic endothelium^18,19^. Single-cell transcriptomic studies have shown that high-fat diet (HFD) alters endothelial gene expression in a tissue- and cell-type-specific manner^20,21^. However, changes in gene expression alone cannot establish how these molecular differences affect signaling dynamics or junction regulation.

Integrating population-specific transcriptional changes into a mechanistic model provides an opportunity to investigate how HFD-associated molecular remodeling may influence VEGF-A responses across distinct lymphatic vascular beds.

Here, we developed a mechanistic computational model of VEGF-A signaling calibrated to experimental data from human dermal LECs and linked downstream RAC1–RhoA dynamics to a predicted junction state (Fig. 1). By incorporating cell-to-cell variability and HFD-associated transcriptional changes, we examined how VEGF-A signaling responses differ across individual cells and LEC populations. The model predicts that VEGF-A dose primarily regulates the duration of zippering-favoring signaling, while HFD-associated molecular changes produce distinct responses across lymphatic vascular beds. These findings provide a quantitative framework for understanding how signaling dynamics and LEC molecular heterogeneity may influence lymphatic junction regulation.

**Figure 1.**
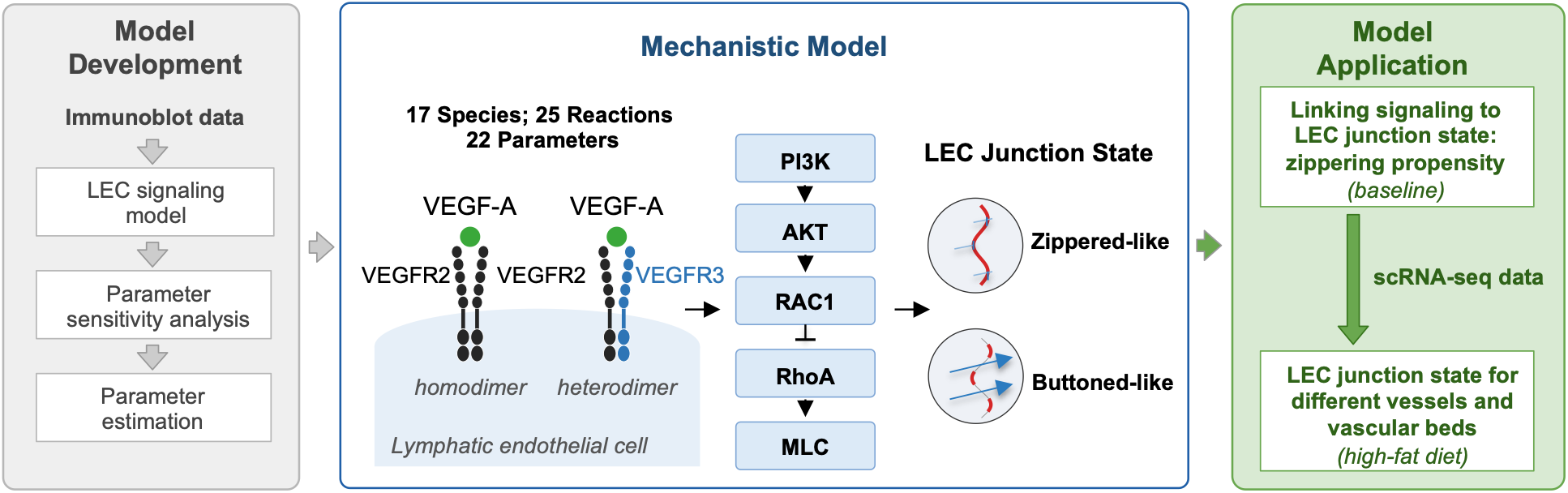
A mechanistic model linking VEGF-A signaling to the junctional state of lymphatic endothelial cells. Overview of model development (left), the model structure (center) and its application (right). **Model Development**: An ordinary differential equation (ODE) model of the VEGF-A signaling in lymphatic endothelial cells (LECs) was calibrated against published immunoblot data^10^ by global sensitivity analysis and parameter estimation. **Mechanistic Model**: VEGF-A activates VEGFR2 homodimers and VEGFR2/VEGFR3 heterodimers, which signal through PI3K and AKT to activate RAC1. RAC1 inhibits RhoA, which drives myosin light chain (MLC) phosphorylation (17 species, 25 reactions, 22 parameters). **Model Application**: The calibrated model was used to predict LEC junction state at baseline and was then combined with single-cell RNA sequencing (scRNA-seq) data from mesenteric and epididymal adipose tissue of mice fed a high-fat diet (HFD) ^20^ to predict junction state under HFD across vessel types and vascular beds.

## RESULTS

### Model training and validation

To capture the signaling events governing LEC junction remodeling, we constructed an ODE model of the VEGF-A–VEGFR signaling cascade, built on pathway architecture established in Human Dermal Lymphatic Endothelial Cells (HDLECs) ^10^. The model represents the core signaling network linking VEGF-A-induced receptor activation to cytoskeletal regulation of junction remodeling, as shown in Supplementary Fig. 1. VEGFR2 homodimers and VEGFR2/VEGFR3 heterodimers form and are activated by VEGF-A, propagating signal through PI3K and AKT1 to RAC1, which non-competitively inhibits RhoA and thereby regulates phosphorylation of MLC (to produce pMLC). Each dimer and downstream signaling node is represented by distinct active and inactive species with separate initial values, where active species catalyze activation of their downstream targets. We assumed that the reactions follow mass-action or Michaelis-Menten kinetics rate laws and constructed the ODE model to describe the rate of change of each species concentrations, based on the reactions it participates in.

To quantitatively fit the ODE model to experimental measurements of VEGF-A-VEGFR signaling in HDLECs, we first used global sensitivity analysis (Supplementary Fig. 2) to identify kinetic parameters and initial species concentrations that most strongly influence the simulated dynamics. We then performed particle swarm optimization (PSO) to estimate the optimal values of these influential parameters by calibrating the model to the experimental data. As a result, we obtained 89 sets of parameters that enable the model to closely reproduce the training data (Supplementary Data 1). The calibrated model reproduced the VEGF-A-VEGFR signaling time course across the pathway, capturing the correct amplitude and temporal ordering of the cascade (R² = 0.972, RMSE = 0.185 fold-change units; Fig. 2a-e). Receptor activation was fast and transient, downstream AKT and RAC1 activation peaked later and persisted longer, and RhoA and pMLC were suppressed over the same window.

**Figure 2.**
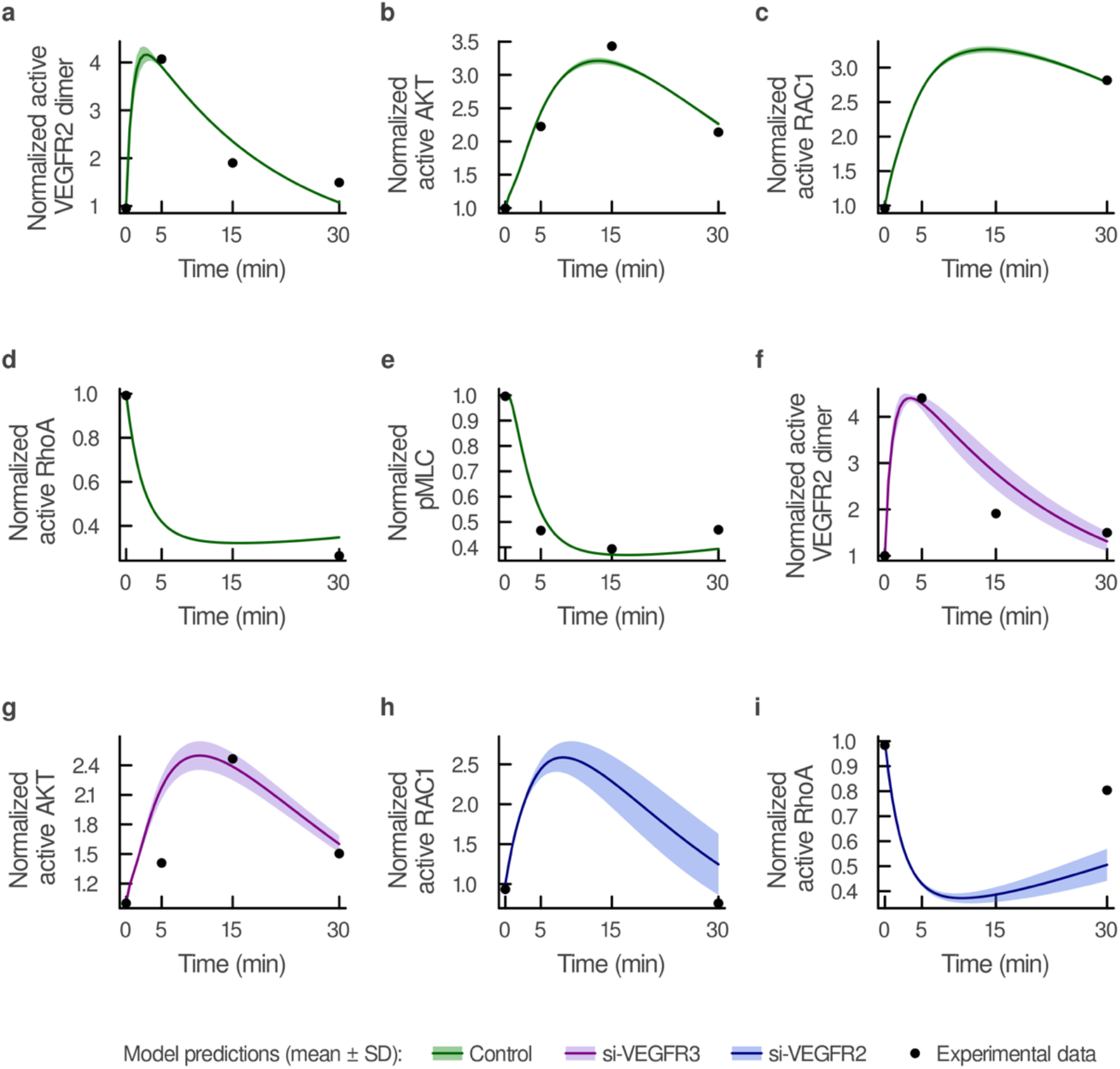
The calibrated model reproduces VEGF-A signaling dynamics in lymphatic endothelial cells and predicts independent receptor-knockdown responses. a–e. Model fits to calibration data^10^ from lymphatic endothelial cells (LECs) stimulated with 50 ng/mL VEGF-A: **a** normalized active VEGFR2 dimer, **b** normalized active AKT **c** normalized pMLC, **d** normalized active RAC1, and **e** normalized active RhoA. **f–i** Model predictions compared with independent data from VEGFR2- or VEGFR3-silenced LECs^10^ that were not used for calibration: **f** normalized active VEGFR2 dimer (si-VEGFR3), **g** normalized active AKT (si-VEGFR3), **h** normalized active RAC1 (si-VEGFR2), and **i** normalized active RhoA (si-VEGFR2). Knockdown was simulated by increasing the degradation rate of the targeted receptor 10-fold without refitting. Lines, mean of the 89 of 100 particle swarm optimization (PSO) parameter sets in which no estimate reached a search bound; shaded bands, ± SD across these 89 sets; dots, mean of n = 4–6 replicate measurements digitized from the original study, expressed as fold change relative to unstimulated control (*t* = 0).

To validate the model, we tested its predictions against an independent set of receptor knockdown measurements that were not used for model fitting. We simulated si-VEGFR2 and si-VEGFR3 knockdown by increasing the corresponding receptors’ degradation rate 10-fold, without re-fitting any other parameters, and compared the resulting dynamics to the knockdown experiments ^10^. We found that model predictions closely matched the experimental data (R² = 0.850, RMSE = 0.381; Fig. 2f–i). The si-VEGFR3 served as a specificity control, damping AKT activation while leaving VEGFR2 activation intact, whereas si-VEGFR2 truncated RAC1 activation and limited the repression of RhoA compared to data obtained in the absence of si- VEGFR2. Together, the calibration and independent knockdown validation establish that the ODE model captures the temporal organization of VEGF-A–VEGFR signaling in LECs and can be used to examine how changes in pathway inputs alter downstream RAC1–RhoA dynamics.

### Quantitatively linking cellular signaling to junction state by defining zippering propensity and response duration

Using the calibrated model, we simulated RAC1/RhoA signaling across four VEGF-A doses (2.5, 10, 50, and 80 ng/mL) (Fig. 3a–b). Simulated trajectories for all species (except VEGF-A) across these doses are shown in Supplementary Fig. 4. Active RAC1 and active RhoA exhibited dose-dependent transient responses but converged toward the same steady state regardless of VEGF-A dose.

**Figure 3.**
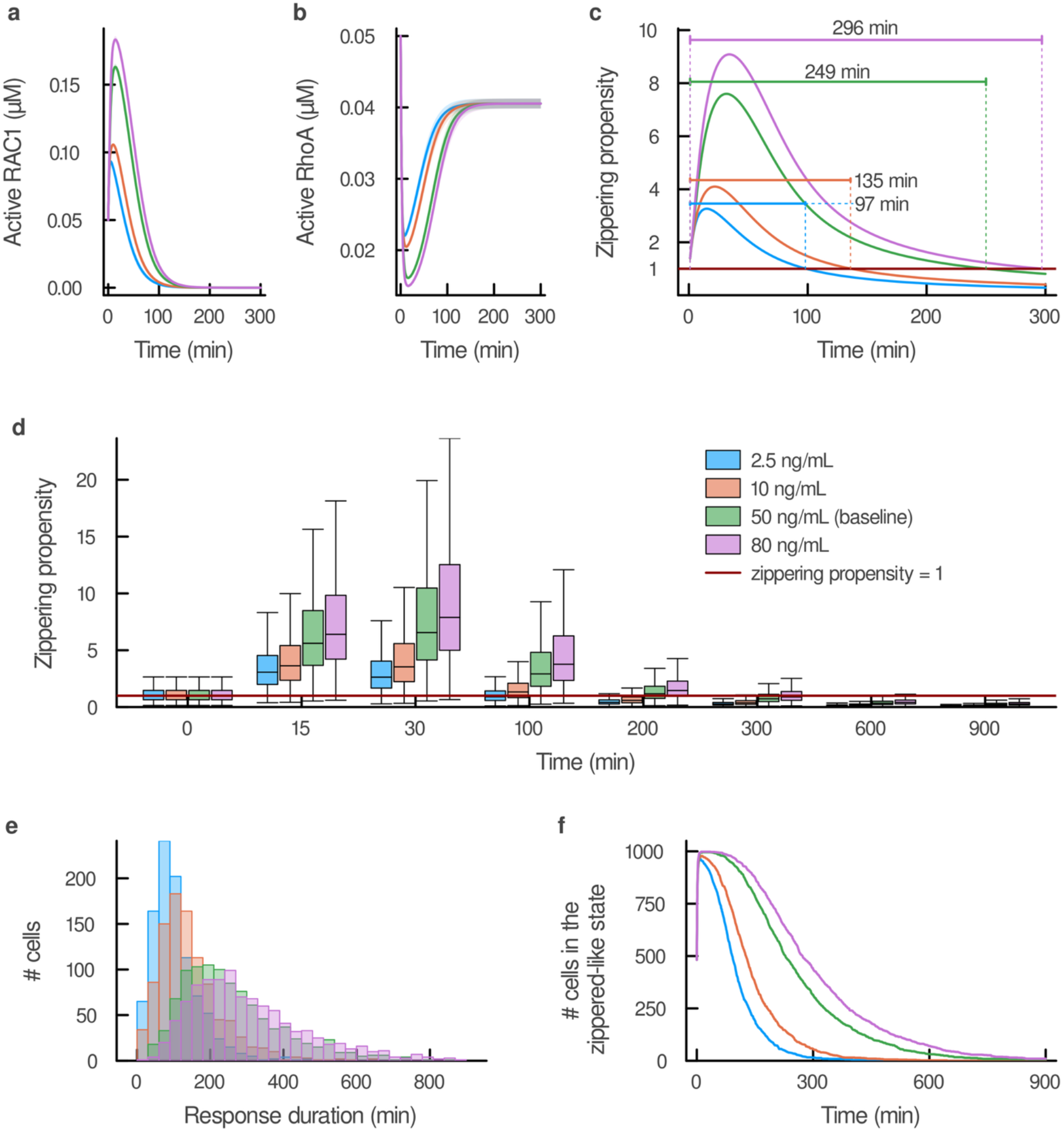
Higher VEGF-A doses increase zippering propensity and prolong response duration in single representative cell and heterogeneous populations. a–c. Single-cell model. **a** Predicted active RAC1 and **b** active RhoA trajectories. **c** Zippering propensity, with the threshold of 1 and the response duration indicated. **d–f** Heterogeneous population model (N = 1,000 simulated cells per dose). **d** Per-cell zippering propensity at eight sampled times (0, 15, 30, 100, 200, 300, 600, and 900 min), shown as grouped box plots. **e** Histograms of response duration. **f** Number of cells in the zippered-like state over 0–900 min. Zippering propensity is the ratio of the areas under the curve (AUC) of active RAC1 and active RhoA; values > 1 denote the zippered-like junction state and values ≤ 1 the buttoned-like junction state. Response duration is the time over which zippering propensity exceeds 1. VEGF-A doses are 2.5 (blue), 10 (orange), 50 (green), and 80 ng/mL (purple). In **a** and **b**, lines show the mean and shaded bands ± SD across the 89 retained particle swarm optimization (PSO) parameter sets; **c–f** use the best-fit parameter set. In **d**, boxes span the interquartile range (IQR), the line marks the median, and whiskers extend to 1.5 × IQR.

Because RAC1 and RhoA exert opposing effects on endothelial junction stability^6,22^, we integrated their activities into a single model-derived metric of junctional behavior. We defined “zippering propensity” as the ratio of cumulative active RAC1 to active RhoA activity, calculated from the area under the curve (AUC) of their simulated trajectories. Values above one represent RAC1-dominant signaling and a predicted zippered-like state, whereas values at or below one represent RhoA-dominant signaling and a predicted buttoned-like state. From the zippering propensity time course, we derived two summary measures: maximum zippering propensity, defined as the peak value of the ratio, and response duration, defined as the cumulative time during which zippering propensity remained above one.

Zippering propensity increased transiently before declining at all VEGF-A doses before declining. Using the best-fit parameter set, response duration increased from 97 min at 2.5 ng/mL to 135 min at 10 ng/mL, 249 min at 50 ng/mL, and 296 min at 80 ng/mL (Fig. 3c). Thus, increasing VEGF-A dose primarily prolonged the RAC1-dominant signaling state, identifying response duration as a key model-derived output of VEGF-A signaling. We next asked whether this dose-response behavior was maintained when cell-to-cell variability in the signaling components was incorporated into the model. To represent a heterogeneous LEC population, we varied the initial concentrations of individual signaling species by sampling from Gamma distributions centered on their baseline model values, with each combination of initial conditions representing an individual simulated cell^23^. We compared populations of 500, 1,000, 10,000, and 20,000 cells and found that a population of 1,000 cells provided stable estimates of zippering propensity while maintaining computational efficiency (Supplementary Fig. 5).

We simulated this heterogenous 1,000-cell population (Supplementary Fig. 6) across the same four VEGF-A doses (2.5, 10, 50, and 80 ng/mL). Although VEGF-A dose had relatively little effect on the initial transition into the predicted zippered-like state, it strongly influenced how long this state persisted and the extent of cell-to-cell variability in response duration (Fig. 3d–f). At the 50 ng/mL VEGF-A dose used for model fitting and calibration, individual simulated cells followed qualitatively similar trajectories, but with different magnitude and duration (supplementary Fig. 7a-c, 50 representative cells). Active RAC1 increased transiently before returning toward baseline, while active RhoA and pMLC initially decreased and subsequently recovered to cell-specific steady states. These differences produced substantial cell-to-cell variability in both the magnitude of zippering propensity and response duration (Supplementary Fig. 7d).

We next examined how VEGF-A dose affected this heterogeneous population over time (Fig. 3d). Higher VEGF-A doses delayed both the timing of peak zippering propensity and the population’s decay into the buttoned-like state, and increased cell-to-cell variability in the response. We analyzed the population-level response across the four doses (Fig. 3d), classifying each cell as in the zippered-like state (zippering propensity > 1) or in the buttoned- like state (zippering propensity ≤ 1). At 2.5 ng/mL, zippering propensity reached its peak magnitude by ∼15 min and then declined, with more than half of cells in the buttoned-like state (median <1) by 100 min. At 10 ng/mL, peak magnitude was reached later, around 30 min, and more than 75% of cells had reached the buttoned-like state (Q3 <1) by 200 min. At 50 ng/mL, approximately 75% of cells were in the buttoned-like state (Q3 <1) by 300 min, and at 80 ng/mL, around half of cells were in the buttoned-like state (median <1) by the same time point.

Consistent with these temporal differences, higher VEGF-A dose also increased the variability of response duration across cells. We plotted the distribution of response duration across the 1,000-cell population at each dose; mean duration nearly tripled across the dose range tested (Fig. 3e). The fraction of cells in a zippered-like junction state rose rapidly and similarly across doses but declined in a strongly dose-dependent manner (Fig. 3f): initially, only 48.2% of cells had a zippering propensity above one, and 95.4–99.8% of cells became zippered-like within 15 min, with only a modest increase with dose. Decay of the fraction of cells in the zippered-like junction state, in contrast, was strongly dose-dependent. In summary, extending the model to a heterogeneous 1,000-cell population revealed that VEGF-A dose acts primarily on the response duration. Nearly all cells are predicted to be in the zippered-like state within minutes across all doses, but the duration of that zippered-like state, its peak timing, and its variability across cells are all scaled with dose.

### Identifying key regulators of junction response dynamics

To identify the signaling features that control junction response dynamics, we performed global sensitivity analysis using response duration and maximum zippering propensity as model outputs. At 50 ng/ml VEGF-A, six kinetic parameters (Vmax, k8, Km, k4, k6, and k5) were identified as influential (Fig. 4a). These parameters primarily regulate the activation and inactivation kinetics of RAC1 and RhoA. For both model outputs, total-order sensitivity indices (ST) exceeded first-order indices (S1), indicating that interactions among parameters contributed substantially to the predicted response.

**Figure 4.**
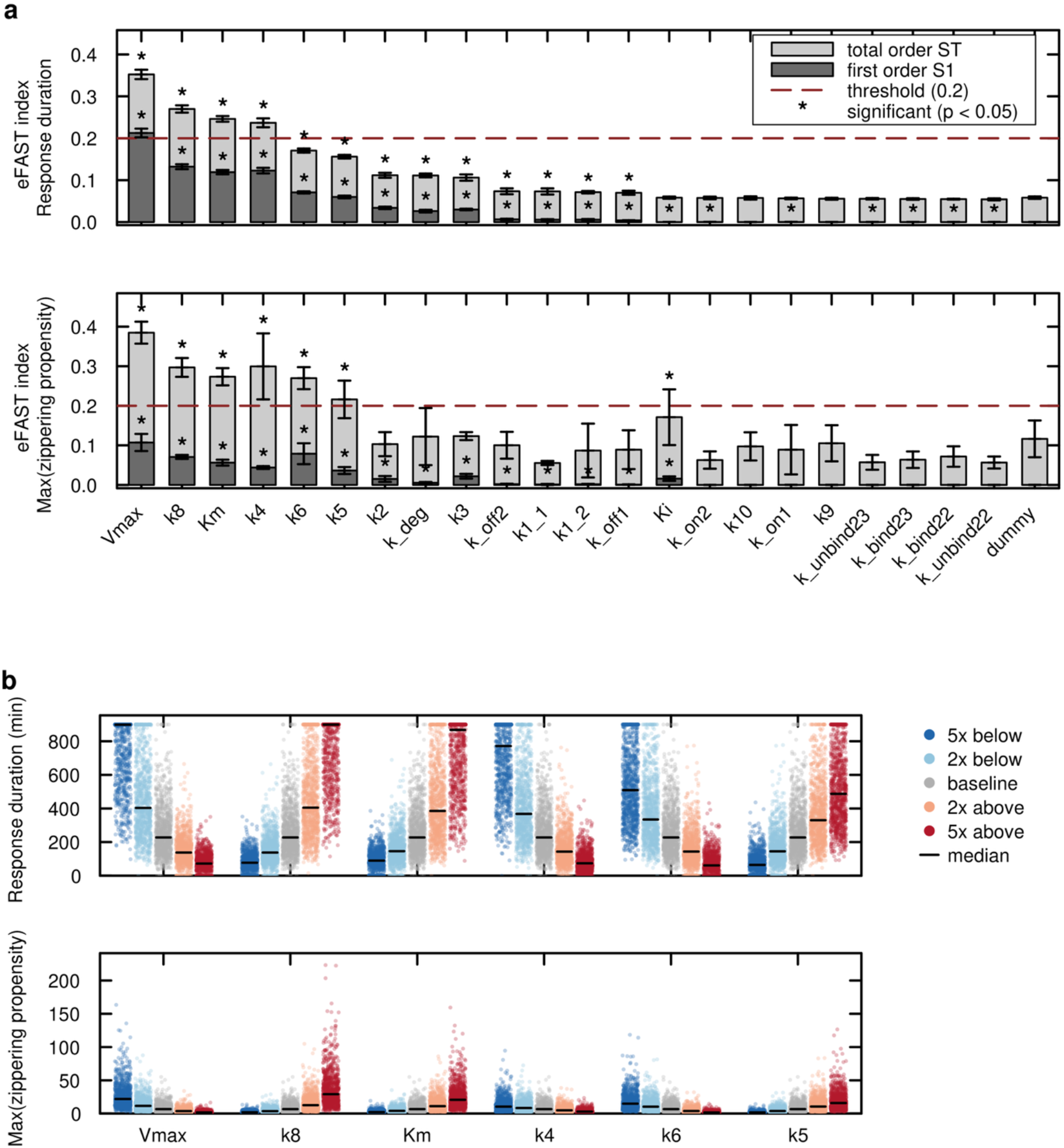
A small set of kinetic parameters controls zippering propensity and response duration in the heterogeneous population model. **a** Global sensitivity analysis by the extended Fourier amplitude sensitivity test (eFAST), with response duration and maximum zippering propensity as outputs. Total-order (ST) and first-order (S1) indices are shown; error bars, SD of 10 eFAST repeats. Asterisks mark parameters whose ST was significantly greater than that of a dummy parameter (one-tailed Welch’s *t*-test, *P* < 0.05). Parameters with ST > 0.2 and a significant difference from the dummy were considered influential: Vmax, maximum rate of RhoA activation; k8, RhoA deactivation; Km, Michaelis constant of RhoA activation; k4, AKT deactivation; k6, RAC1 deactivation; and k5, RAC1 activation. **b** Effect of scaling each influential parameter to 0.2-, 0.5-, 2-, and 5-fold its best-fit value, with all other parameters held at baseline. Each point is one simulated cell (N = 1,000 cells); the median is marked. Simulations ran for 900 min; some cells remained in the zippered-like state at 900 min.

We next systematically perturbed each influential parameter to determine how it affected the heterogeneous LEC population. Each of the six influential parameters was independently scaled to 0.2, 0.5, 1, 2 and 5-fold of its best-fit value, with all others held at baseline, and the model was re-run for every cell of the same gamma distribution-sampled population of N = 1,000 virtual HDLECs used in the results presented above (Fig. 4b). Increasing maximum rate of RhoA activation (Vmax), AKT deactivation (k4) or RAC1 deactivation (k6) shortens response duration and weakens zippering propensity; increasing RhoA deactivation (k8), the Michaelis constant of RhoA activation (Km) or RAC1 activation (k5) prolongs and strengthens the zippering propensity. Under some extremes, including reduced Vmax and increased k8, a large fraction of cells reached the 900-minute simulation endpoint still in the zippered-like state. Thus, the response duration in these conditions represent lower bound estimates rather than a true window length.

Global sensitivity analysis of the model’s initial conditions identified the initial RAC1, RhoA and active RhoA abundance as major determinants of the population response (Supplementary Fig. 8). Across the heterogeneous population, response duration correlated positively with RAC1 (Pearson r = 0.58, Spearman ρ = 0.59) and negatively with RhoA (r = −0.51, ρ = −0.53) (Supplementary Fig. 9). Maximum zippering propensity showed a stronger positive correlation with RAC1 (r = 0.69, ρ = 0.71) and a weaker negative correlation with RhoA (r = −0.37, ρ = −0.42). Together, the sensitivity analyses identified key RAC1–RhoA antagonism parameters that regulate the population-level response.

### High-fat diet remodels LEC composition and VEGF signaling genes in a depot- and subtype-specific manner

Having identified signaling features that control the predicted junction response, we next asked whether changes in LEC molecular state could reshape these dynamics in vivo. We used diet-induced obesity as a biologically relevant context because HFD remodels the lymphatic endothelium and provides an opportunity to examine whether transcriptional changes across distinct lymphatic beds and vessel types translate into different predicted signaling behaviors.

To define HFD-associated LEC states, we analyzed published single-cell RNA-sequencing data from murine mesenteric and epididymal adipose tissue after 12 weeks of normal chow or HFD feeding^20^. The dataset included 10,785 endothelial cells from mesenteric adipose tissue and 13,321 from epididymal adipose tissue across dietary conditions. LECs represented approximately 10–15% of the endothelial compartment in both depots and diets (Fig. 5a-b and Supplemental Fig. 10). Sub-clustering identified four LEC populations based on previously described markers^24^ (Fig. 5c). We found that HFD altered LEC composition differently between the two depots (Fig. 5d). In mesenteric adipose tissue, capillary LECs increased from 44.8% to 61.6%, whereas collector LECs decreased from 24.0% to 14.3%. In epididymal adipose tissue, capillary LECs decreased from 69.2% to 58.3%, whereas collector LEC abundance was relatively unchanged (13.4% versus 15.5%). Pre-collector LECs increased in both depots, whereas proliferative LECs decreased. Differential expression analysis further revealed depot- and subtype-specific responses to HFD (Fig. 5e). Within the VEGF-A–VEGFR signaling network, mesenteric capillary LECs showed reduced expression of the VEGF receptors *Kdr* and *Flt4* together with increased *Rac1* and *Rhoa*. These changes were restricted to the capillary population, with no pathway genes meeting the significance threshold in mesenteric pre-collector or collector LECs. In epididymal adipose tissue, capillary LECs instead showed increased *Pik3ca* expression without significant changes in other genes in this pathway.

**Figure 5.**
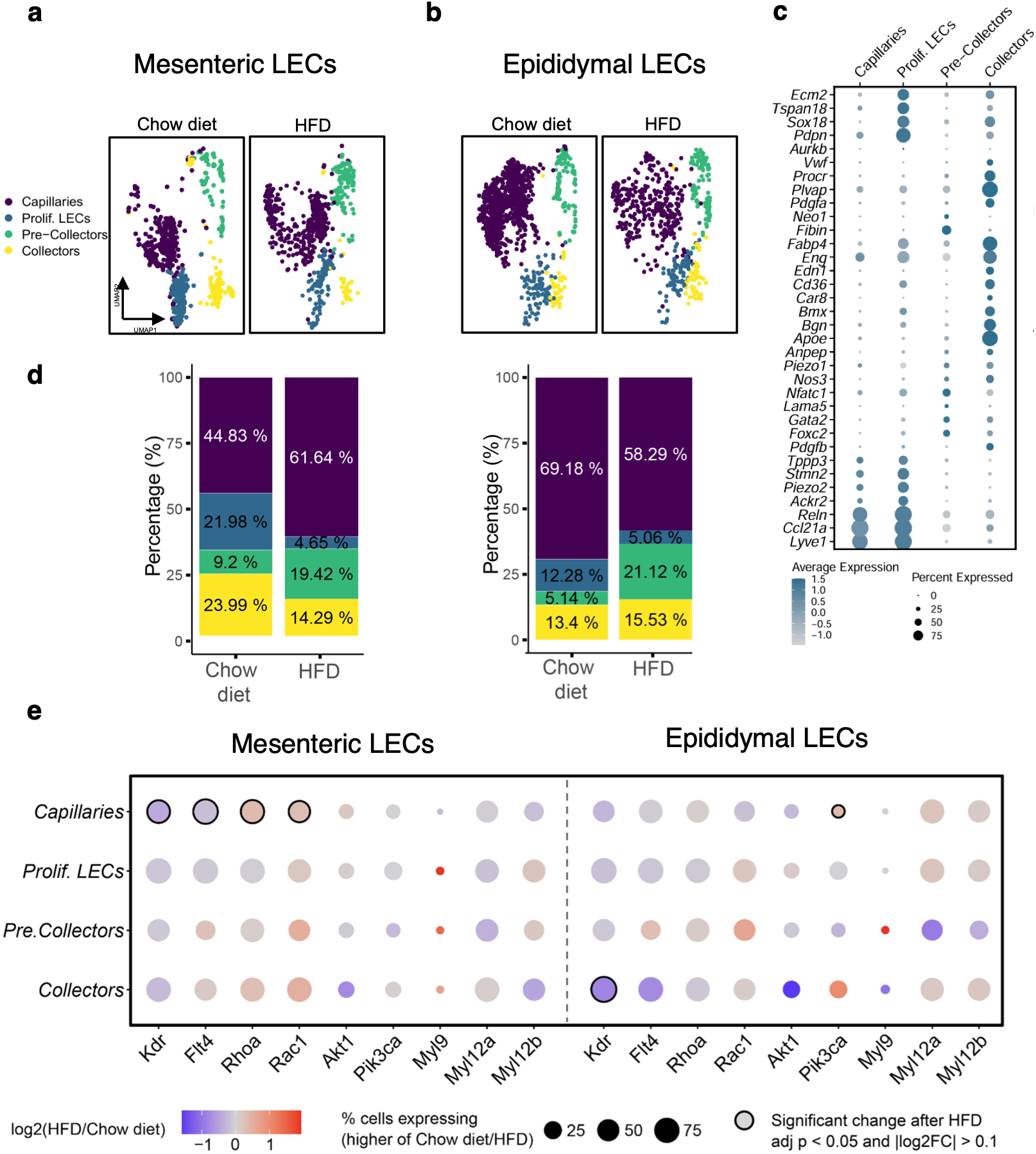
High-fat diet remodels LEC composition and VEGF signaling genes in a depot- and subtype-specific manner. Single-cell RNA sequencing data from mesenteric and epididymal adipose tissue of mice fed normal chow or a high-fat diet (HFD) for 12 weeks^20^ were reanalyzed. **a**, **b** Uniform manifold approximation and projection (UMAP) of lymphatic endothelial cells (LECs) from **a** mesenteric tissue (chow, 696 cells; HFD, 623 cells) and **b** epididymal tissue (chow, 1,114 cells; HFD, 587 cells), comparing chow and HFD. **c** Dot plot of marker genes used to annotate LEC subpopulations (capillaries, proliferating LECs, pre- collectors, and collectors). Dot size indicates the percentage of cells in each subpopulation expressing the gene; dot color indicates scaled average expression. **d** Composition of mesenteric (left) and epididymal (right) LECs on chow and HFD. **e** Fold change in expression between HFD and chow within each depot for genes passing significance filtering (two-sided Wilcoxon rank-sum test, Benjamini–Hochberg adjusted *P* < 0.05, |log2 fold change| > 0.1). Dot size indicates the percentage of cells expressing the gene (the larger of the chow and HFD values); color indicates log2 fold change (red, upregulated in HFD; blue, downregulated in HFD).

Epididymal collector LECs showed reduced *Kdr* expression. Thus, HFD altered components of the VEGF-A signaling network in patterns that depended on both anatomical depot and LEC subtype.

### Diet-induced obesity models and strategies to restore baseline behaviors

Because HFD produced distinct changes in VEGF-A signaling components across LEC populations, we next asked whether these molecular states were sufficient to alter predicted junction response dynamics. We generated three subpopulation-specific models representing mesenteric capillary, epididymal capillary, and epididymal collector LECs (**Table 1**) by perturbing the initial conditions and/or kinetic parameters implicated by each subpopulation’s HFD gene-expression changes. We considered these altered initial conditions and parameters across a range of magnitudes rather than translating mRNA fold-changes directly into specific parameter values (see Methods). HFD-associated perturbations produced distinct effects on predicted response duration across the three LEC populations (Fig. 6a-c). Mesenteric capillary LECs and epididymal collector LECs models shifted toward shorter response durations, whereas the epididymal capillary LECs model shifted toward longer response durations. These population-specific effects were maintained across the range of VEGF-A doses tested, indicating that HFD-associated molecular remodeling can alter the temporal response to VEGF- A differently depending on LEC subtype and anatomical depot.

**Figure 6.**
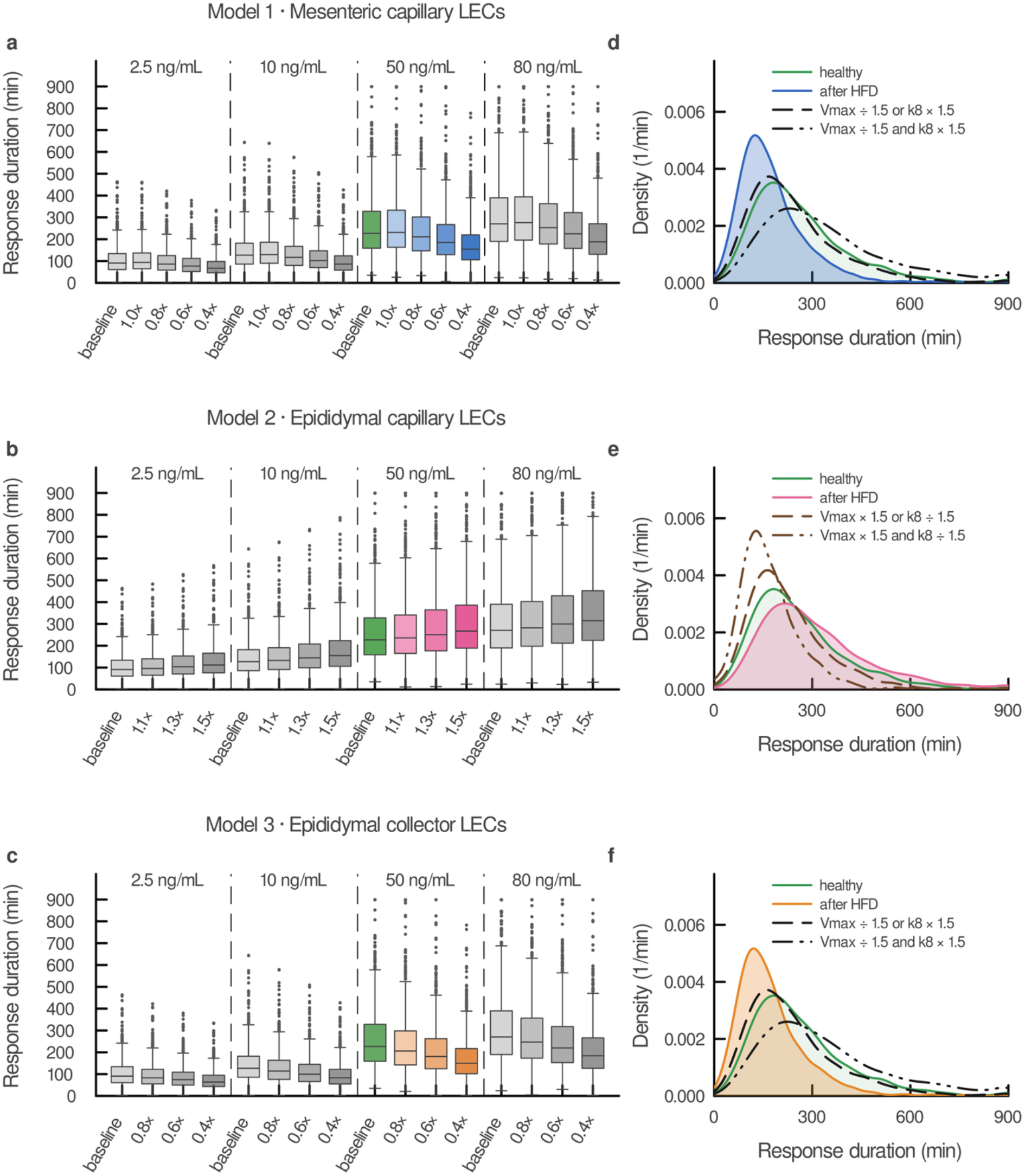
High-fat diet alters response duration in three subpopulation-specific models, and tuning RAC1–RhoA antagonism restores baseline in simulations. **a** Response duration at four VEGF-A doses (2.5, 10, 50, and 80 ng/mL) across increasing perturbation magnitude for **a** Model 1, mesenteric capillary lymphatic endothelial cells (LECs); **b** Model 2, epididymal capillary LECs; and **c** Model 3, epididymal collector LECs, with perturbations as defined in Table 1. Within each dose, the leftmost box is the unperturbed model and successive boxes show increasing perturbation magnitude (darker shading). The 50 ng/mL group is colored (green, baseline; blue, Model 1; pink, Model 2; orange, Model 3); other doses are gray. Boxes span the interquartile range (IQR), the line marks the median, whiskers extend to 1.5 × IQR, and points beyond are outliers. **d–f** Distributions of response duration at 50 ng/mL VEGF-A for **d** Model 1, **e** Model 2, and **f** Model 3, comparing the healthy baseline (green), the most strongly perturbed high-fat diet model (filled; blue, pink, or orange), and three simulated treatments targeting the RAC1–RhoA axis: Vmax (maximum rate of RhoA activation) or k8 (RhoA deactivation) changed 1.5-fold individually, or both together, in the direction that restores baseline (Models 1 and 3: Vmax ÷ 1.5 and k8 × 1.5, black; Model 2: Vmax × 1.5 and k8 ÷ 1.5, brown). N = 1,000 simulated cells per condition.

**Table 1.**
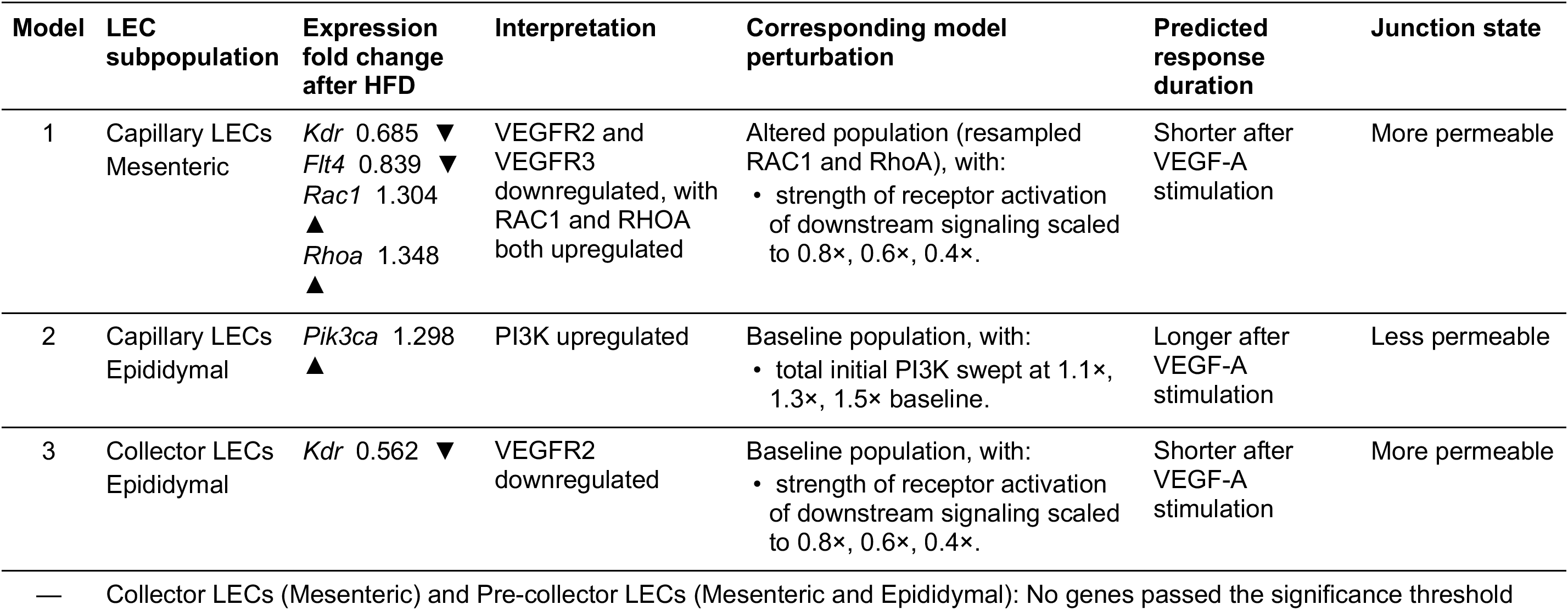
LEC subpopulation phenotypes after high-fat diet and their corresponding model perturbations. For each lymphatic endothelial cell subpopulation, differentially expressed genes after high-fat diet (fold-change relative to chow; ▴ upregulated, ▾ downregulated) were translated into a corresponding perturbation of the computational model. Predicted response duration after VEGF-A stimulation and the resulting junction state are model outputs.

Using the LEC subtype- and depot-specific mathematical models of VEGF-A-mediated signaling altered by HFD, we tested whether the response could be rescued by tuning Vmax and k8, the two RAC1–RhoA parameters shown by sensitivity analyses to influence junction response. Specifically, we next asked whether modifying this signaling axis could shift the HFD-associated responses toward baseline. In the mesenteric capillary and epididymal collector model, simulations show that decreasing Vmax (maximum rate of RhoA activation) or increasing k8 (RhoA deactivation) 1.5-fold individually shifted the shortened distributions for response duration back toward baseline (healthy condition) (Fig. 6d–f). Similarly, increasing Vmax (maximum rate of RhoA activation) or decreasing k8 (RhoA deactivation) restored the healthy condition for the model representing epididymal capillary LECs. In addition, altering both parameters together overshot the baseline response in the corrective direction. These simulations predict that HFD-associated molecular remodeling produces heterogeneous, bed- and vessel-type-specific junction outcomes, yet these divergent responses remain sensitive to modulation of a common RAC1–RhoA regulatory axis. Thus, Vmax (maximum rate of RhoA activation) and k8 (RhoA deactivation) emerge as a shared control point capable of shifting HFD-altered, population-specific responses back toward baseline, although the direction of the required adjustment depends on the bed and vessel type.

## DISCUSSION

We developed a computational mechanistic model that links VEGF-A–VEGFR signaling to a predicted junction state in heterogeneous populations of LECs. The model makes three main predictions. First, across VEGF-A doses, RAC1-dominant signaling begins at a similar time, but its peak and especially its duration increase with dose. Second, HFD-associated transcriptional changes shift response duration in opposite directions in different LEC subpopulations. Third, RAC1–RhoA regulation acts as a shared control point: adjusting it can move all three HFD- affected populations back toward baseline, although the required direction differs between populations. Together, these findings provide a quantitative framework for how signaling dynamics and LEC molecular state combine to regulate lymphatic junctions.

Our model builds on experimental evidence that VEGF-A through VEGFR2/VEGFR3, PI3K, AKT and RAC1 to regulate LEC junction organization and lymphatic permeability^10^. While these studies established the molecular pathway linking VEGF-A stimulation to junction remodeling, how signaling dynamics influence this response remains poorly understood. Our simulations suggest that VEGF-A dose primarily determines the duration of RAC1-dominant signaling rather than the initial transition toward a zippering-favoring state. Across the concentrations tested, nearly all simulated cells rapidly developed RAC1-dominant signaling, whereas higher VEGF-A concentrations prolonged this response and increased variability in its duration. These findings suggest that the persistence of signaling, rather than its initial activation alone, may be important for lymphatic junction regulation. Because junction remodeling requires coordinated changes in cytoskeletal organization and cell–cell adhesion, differences in signaling duration could influence how long endothelial junctions remain in a zippering-favoring state. This prediction highlights the importance of examining junction organization over time rather than relying on measurements at individual endpoints.

Mechanistic VEGF models have been developed mainly for blood endothelial cells^15^, describing ligand–receptor binding, receptor dimerization and downstream PI3K–AKT and ERK signaling, primarily in the context of angiogenesis, proliferation and migration^25^. Computational work on the lymphatic system has instead focused on fluid transport and lymphangion pumping^16^. Our model differs from these in three ways: (i) it is built for and calibrated to LEC data, and it describes VEGF-A signaling in LECs, including VEGFR2 homodimerization and VEGFR2/VEGFR3 heterodimerization, rather than the canonical VEGF-C/VEGFR3 pathway primarily studied in lymphatics (ii) it extends signaling to the balance between RAC1 and RhoA, which allows receptor-level events to be linked to a predicted junction state; and (iii) it represents cell-to-cell variability in protein abundance, rather than a single average cell. To our knowledge, this is the first mechanistic model of VEGF-A signaling designed to predict junction behavior in lymphatic endothelium.

Cell-to-cell heterogeneity was an important feature of the model. Endothelial cells in the same population can differ substantially in the abundance of receptors and signaling proteins, which can affect both the strength and the duration of pathway activation^23^. In our simulations, nearly all cells quickly shifted toward a predicted zippered-like state at every VEGF-A dose, but how long they stayed in this state varied widely between cells, and this variation increased at higher doses. Population averages may therefore hide differences in signaling persistence that matters for junction behavior. This may be especially relevant in lymphatic vessels, where permeability also depends on the combined behavior of neighboring cells (e.g. macrophages, dendritic cells, etc.). Differences in signaling duration may contribute to spatial variation in junction remodeling, although this possibility remains to be tested experimentally.

LEC molecular state added a second layer of heterogeneity. When we incorporated HFD-associated transcriptional changes, the same VEGF-A signaling network produced different responses depending on vascular bed and vessel type. Response duration shifted in opposite directions in mesenteric and epididymal capillary LECs, and between capillary and collector LECs within the epididymal bed. These results extend previous reports that HFD remodels endothelial transcriptional programs in a population-specific way^20^ by suggesting that these molecular differences translate into distinct signaling dynamics. Importantly, these outcomes could not have been predicted from gene expression changes alone. For example, mesenteric capillary LECs upregulated *Rac1*, which on its own would be expected to prolong zippering. However, the model predicted shorter responses, because reduced *Kdr* and *Flt4* and increased *Rhoa* outweighed this effect. Thus, the impact of changing any single component depends on its interactions within the wider network. These findings are relevant to the growing evidence linking lymphatic dysfunction with obesity and metabolic disease^26^. Experimentally altering lymphatic junctions changes dietary lipid transport and susceptibility to diet-induced obesity^10^, and obesity in turn causes structural and functional lymphatic abnormalities.

Mesenteric lymphatic leakiness and remodeling are associated with visceral fat expansion and impaired metabolic health^27^, and improving mesenteric lymphatic function can reduce these abnormalities^19^. Our prediction that HFD shortens zippering-favored signaling in mesenteric capillary LECs is consistent with this increased mesenteric permeability. Fang and colleagues highlighted the need to combine VEGF–VEGFR modeling with tissue-specific data to understand vascular responses in obesity^28^. Our study applies this approach to lymphatic endothelium and suggests that HFD does not affect the lymphatic vasculature uniformly but instead alters VEGF-A response dynamics in a bed- and vessel-specific manner.

The physiological implications of these predictions likely depend on the anatomical location and functional specialization of lymphatic vessels. Initial lymphatic capillaries typically exhibit discontinuous button-like junctions that facilitate fluid and macromolecule uptake, whereas collecting vessels maintain continuous zipper-like junctions that support lymph retention during transport^2^. In mesenteric capillary LECs, HFD-associated molecular changes shortened the predicted duration of zippering-favoring signaling, potentially favoring a more permissive junctional state. In contrast, epididymal capillary LECs exhibited prolonged zippering-favoring signaling, which could restrict fluid and macromolecule uptake from adipose tissue. Epididymal collector LECs showed shorter responses, potentially compromising the maintenance of continuous junctions required for lymph retention. These predictions are consistent with evidence that obesity alters lymphatic permeability and transport^19,27^, although the model does not directly establish changes in vessel function. Importantly, the opposite responses predicted for capillary LECs from different adipose depots suggest that HFD-associated lymphatic remodeling cannot be explained by a uniform change in VEGF-A signaling. Instead, the consequences of metabolic stress may depend on the molecular state and functional specialization of individual lymphatic endothelial populations.

Despite the divergent responses to HFD-associated molecular changes, sensitivity analyses identified common determinants of junction-associated signaling dynamics. The rates of RhoA activation and inactivation, together with the relative amounts of RAC1 and RhoA, strongly influenced both zippering propensity and response duration. Adjusting Vmax (maximum rate of RhoA activation) or k8 (RhoA deactivation) shifted the HFD response-duration distributions toward baseline in all three populations. However, the direction of the required adjustment differed between populations. The model therefore does not predict that uniformly increasing or decreasing RAC1–RhoA signaling would normalize lymphatic responses across tissues. Instead, one regulatory module can produce different outcomes depending on the molecular state of the cell. This context dependence may be important therapeutically, as targeting Rho/ROCK or RAC1 signaling could have different effects across lymphatic beds.

Strategies to modify lymphatic permeability may therefore need to consider tissue and vessel type. Our study has several limitations. First, the calibration and validation data end at 30 minutes, so the later dynamics that determine response duration (up to 900 minutes) are model predictions and require experimental confirmation. Related, changing the simulation time would affect the assumption to neglect protein production. Second, the model was calibrated on human dermal LECs and then applied to mouse mesenteric and epididymal LEC subpopulations, which may differ in signaling. Third, the model considers only VEGF-A–VEGFR signaling. It does not include VEGF-C, the main lymphatic ligand for VEGFR3, or other signaling and mechanical inputs that regulate LEC junctions^7^. Fourth, the degree of cell-to-cell variability and the threshold used to classify zippered-like and buttoned-like states are modeling choices. Finally, zippering propensity is derived from RAC1 and RhoA activity and has not yet been calibrated against direct measurements of junction structure. Measuring VE-cadherin organization and permeability under matched conditions will be important to test these predictions. In addition, the HFD models were based on changes in mRNA rather than protein. The scRNA-seq dataset is valuable because it resolves molecular changes across lymphatic subpopulations, since LECs were specifically enriched, by design which is difficult with protein-based methods. However, mRNA levels do not always predict protein abundance or activity^29^. We therefore modeled HFD-associated changes across a range of perturbation sizes rather than converting transcript fold changes directly into model parameters. These simulations should be viewed as predictions of how HFD-associated molecular states may alter signaling dynamics, not as direct measurements of pathway activity.

In summary, this study presents, to our knowledge, the first mechanistic signaling model of lymphatic endothelium. It identifies response duration as a key feature of VEGF-A signaling in LECs and predicts that HFD-associated molecular remodeling alters this temporal response across lymphatic beds and vessel types. Despite these divergent responses, RAC1–RhoA regulation (in particular, activation and deactivation of RhoA) remains a shared control point that can shift each population toward baseline. These findings highlight signaling duration as an important dimension of lymphatic junction regulation and provide testable predictions for how metabolic stress affects different parts of the lymphatic vasculature.

## METHODS

### Mathematical model

We constructed a computational model of VEGF-A-VEGFR signaling in lymphatic endothelial cells (LECs). A system of nonlinear ordinary differential equations (ODEs) was formulated to describe the rate of change of the species’ concentration. The model is composed of 17 ODEs to predict the concentrations of the 17 species in the signaling network over time.

The model file is provided in the Supplementary Note 1, and kinetics parameters and initial values are in Supplementary Table 1-2. Each signaling intermediate that exists in two conformational or nucleotide-bound states is represented as two distinct species, denoted "active" when phosphorylated or GTP-bound.

The model species divide into membrane receptors and intracellular downstream signaling species. For the membrane receptors, we used prior literature^30^ to set the ranges of receptor dimerization rates, ligand-binding rates, and receptor concentrations. For the downstream species, we drew nominal values and set ranges for sensitivity analysis from prior publications^11,31^ before calibrating against experimental data. The baseline VEGF-A stimulus was fixed at 1.32 × 10⁻³ µM, corresponding to 50 ng/mL of a 38 kDa ligand^32^.

The steady state of the model does not depend on VEGF-A dose. All species converge to the same steady state with or without VEGF-A stimulation, because steady-state values are constrained by receptor availability. We neglect receptor production, as it generally occurs on a longer time scale than what we simulate with the model and in response to gene expression changes. We therefore focused our analyses on the transient dynamics that follow VEGF-A stimulation.

### Extracting experimental measures of species in the signaling pathway

Experimental measurements were extracted from the published bar graphs using PlotDigitizer (https://plotdigitizer.com/app), and the individual replicate data points (n = 4–6 per condition) were digitized. Replicate values were then averaged to obtain the mean response for each species and time point. In the original study, all measurements were normalized to the unstimulated control (t = 0), so the extracted values represent fold change relative to this baseline. The extracted data are provided in Supplementary Table 3.

### Model outputs: zippering propensity and response duration

To summarize the balance between signaling that promotes junction zippering and signaling that promotes junction buttoning as a single time-resolved quantity, we defined the zippering propensity, Z(t), as the ratio of the time-integrated activities of active RAC1 and active RhoA:

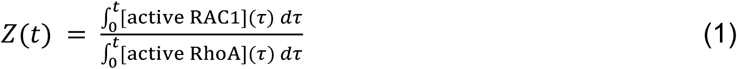

Because both integrals vanish at t = 0, Z(0) was defined as the limit of equation (1) as t → 0, which equals the ratio of the pre-stimulation active concentrations, Z(0) = initial active RAC1/initial active RhoA. Z(t) > 1 indicates RAC1-dominant signaling, corresponding to a zippered-like state, whereas Z(t) ≤ 1 indicates RhoA-dominant signaling, corresponding to a buttoned-like state. We used cumulative rather than instantaneous activities because junctional remodeling reflects signaling integrated over time rather than instantaneous activity. Integrals were computed by trapezoidal integration of the solver output on a uniform grid (Δt = 1 min).

The response duration was defined as the cumulative time (in minutes) during which Z(t) > 1.

### Global sensitivity analysis

There are limited quantitative experimental measurements available to specify the values of the kinetic parameters and initial conditions. We therefore used global sensitivity analysis to identify the most influential model inputs and reduce the number of parameters to be estimated, applying the extended Fourier amplitude sensitivity test (eFAST)^33^, as in our previous work^34^. eFAST yields two indices for each input: the first-order index (*S1*), the fraction of output variance explained by variation in that input alone, and the total-order index (*ST*), which additionally captures the input’s higher-order interactions with all other inputs.

All inputs were sampled uniformly between literature-derived lower and upper bounds, using 3,000 samples per search curve and 10 repeats. The outputs were the concentrations of each of the 17 species at *t* = 5, 15, and 30 min. *ST* was averaged over the 10 resampling runs and compared with the dummy *ST* using a one-tailed Welch’s t-test (α = 0.05). (Supplementary Fig. 2).

After calibration, a second eFAST analysis was performed to identify the kinetic parameters and initial conditions that most strongly influence the model predictions. In this analysis, the outputs were maximum zippering propensity and response duration, and each input was sampled from one order of magnitude below to one order of magnitude above its fitted value. An input was retained as influential if its *ST* exceeded 0.2 and was significantly greater than the dummy in at least one output.

### Parameter estimation

The equilibrium dissociation constant of VEGF-A binding to its receptors is well characterized, whereas the individual association and dissociation rate constants are not. We therefore fixed *K*D = 1.5 × 10^−4^ µM^30^ and computed the association rate constants from the estimated dissociation rate constants (*kon = koff/KD*); this constraint was enforced within the objective function. Together, the sensitivity screen and the affinity constraint reduced the 38 candidate inputs (22 kinetic parameters and 16 initial conditions) to eight influential inputs, which we curated manually to be weakly correlated with one another: six kinetic parameters (*k_deg, k_off1, k_off2, k4, k9, Vmax*) and two initial conditions (*PI3K, RAC1*).

These eight inputs were estimated by fitting the model to the experimental data using particle swarm optimization (PSO)^35^, with the same bounds as in the eFAST analysis. PSO is a population-based, derivative-free optimization method inspired by the collective movement of a flock of birds searching for food. A population of candidate parameter sets (“particles”) moves through parameter space; the objective function is evaluated at each particle’s position, and each particle updates its trajectory using both its own best position and the best position found by the swarm, so that the population progressively converges toward parameter sets that minimize the objective function.

The objective function was the negative log-likelihood (NLL) under an independent Gaussian error model, with both model predictions and observations normalized to their *t* = 0 values:

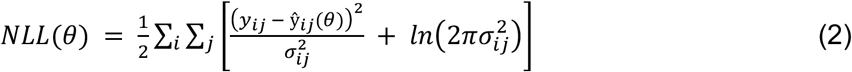

where *y*ij and σij are the measured mean and error of species *i* at time point *j*, and *ŷ*ij(θ) is the corresponding model prediction for parameter vector θ. We used the ParticleSwarm algorithm implemented in Optim.jl with 80 particles and 100 iterations. The optimization was run independently 100 times from the same initial point, using a different random seed for each run (seed = 20250910 + run − 1). The estimates from all 100 runs, ranked by objective function value, are given in Supplementary Data 1, parameter distributions shown in Supplementary Fig. 3. Of these 100 runs, we retained the 89 parameter sets in which no parameter estimate reached its lower or upper bound.

### Population-level model

To account for cell-to-cell heterogeneity, we simulated a population of virtual LECs in which the initial concentration of each species was sampled independently from a Gamma distribution. The Gamma distribution was parameterized by a shape parameter, *a*, and a scale parameter, *b*, which are related to the mean, *m*, and standard deviation, *sd*, by 1/*a* = *sd*^2^/*m*^2^ and *b* = *sd*^2^/*m* (so that *a*·*b* = *m*). For each species, *m* was set to the baseline initial concentration, and *a* was fixed at 5.5 (coefficient of variation ≈ 0.43) based on^36^, as in our previous work^37^.

To select the population size, we compared simulations of 500, 1,000, 10,000, and 20,000 independently sampled cells, which produced similar distributions of zippering propensity over time (Supplementary Fig. 5a). Using the 20,000-cell simulation as the reference, we computed the maximum deviation of the median, first quartile (Q1), and third quartile (Q3) of zippering propensity at *t* = 0, 15, 30, 100, 200, 300, 600, and 900 min for each VEGF-A dose (Supplementary Fig. 5b). Increasing the population from 500 to 1,000 cells reduced the maximum deviation more than increasing it from 1,000 to 10,000 cells, and all maximum deviations for the 1,000-cell population were below 5%. Given the substantially greater computational cost of the larger populations, we used 1,000 cells for all population-level analyses.

Population responses were simulated at VEGF-A concentrations of 2.5, 10, 50 (baseline), and 80 ng/mL. The 1,000-cell population was generated once and reused across all doses and conditions, so that comparisons are paired cell by cell (initial conditions for all 1,000 cells are listed in Supplementary Data 2).

### Single cell RNA-seq data analysis

We analyzed a publicly available single-cell RNA-seq dataset of mesenteric and epididymal adipose tissue from mice fed a high-fat diet (HFD) or normal chow (NC)^20^ (GEO accession GSE235192). LECs were identified using previously described marker genes^24^. Differential expression between HFD and NC was assessed separately for each LEC subpopulation (mesenteric capillary, epididymal capillary, and epididymal collecting LECs) using the FindMarkers function in Seurat v5.4.0^38^ in R 4.5.2, with the default Wilcoxon rank-sum test on log-normalized expression.

The analysis was restricted to a curated panel of VEGF-A–VEGFR pathway genes corresponding to model species: *Kdr* (VEGFR2), *Flt4* (VEGFR3), *Pik3ca* (PI3K), *Akt1* (AKT), *Rac1* (RAC1), *Rhoa* (RhoA), *Myl9* (MLC)*, Myl12a* (MLC)*, and Myl12b* (MLC). For each gene, we extracted the mean log-normalized expression in each condition, together with the average log2 fold change (HFD vs. NC) and its adjusted *p*-value. *P*-values were adjusted using the Benjamini–Hochberg method, and genes with adjusted *p* < 0.05 and |log2 fold change| > 0.1 were considered differentially expressed.

### Model perturbations informed by HFD gene expression change

We translated the HFD-associated expression changes in each LEC subpopulation into model perturbations using two strategies: resampling the initial conditions of the cell population or sweeping the corresponding kinetic parameters across a physiologically plausible range (Table 1).

Each subpopulation’s differentially expressed genes were mapped onto model components as follows. Downregulation of the VEGF receptors *Kdr* (VEGFR2) and *Flt4* (VEGFR3) was represented as reduced strength of receptor-driven downstream activation (*k1_1* and *k1_2*, respectively). Upregulation of *Pik3ca* was represented as an enlarged initial PI3K pool. Changes in *Rac1* and *Rhoa* were represented by resampling total RAC1 and total RhoA (Supplementary Fig. 11): for these two, the mean of the Gamma resampling distribution was scaled directly by the corresponding gene-expression fold change with cell-to-cell variability still supplied by the Gamma sampling, so each perturbed condition remains a distribution rather than a point. All other species were held at their baseline resampling distributions.

### Numerical methods and software

All models were implemented in Julia 1.10.6 using Catalyst.jl^39^, ModelingToolkit.jl^40^, DifferentialEquations.jl^41^, Optim.jl^42^, Distributions.jl^43^, and GlobalSensitivity.jl^44^ within a single project environment; exact package versions are pinned in the Project.toml and Manifest.toml files deposited with the code. Model calibration and all other simulations used Tsit5, an explicit fifth-order Runge–Kutta method, with relative and absolute tolerances of 10^−6^. All sensitivity analyses used Rodas5, a stiff, A- and L-stable Rosenbrock solver, with the same tolerances, because the sampled parameter ranges span several orders of magnitude and include stiff regions of parameter space. Calibration was performed on a SLURM-managed computing cluster using a single node with 8 CPUs and 16 GB of RAM.

### Statistics and Reproducibility

All statistical analyses were performed using publicly available software as detailed in the Methods and Figure legends. Calibration and validation data were digitized from published measurements (n = 4–6 biological replicates per condition), and fit was assessed by R² and RMSE. Predictions are shown as the mean ± SD across the 89 parameter sets from 100 particle swarm optimization runs. Sensitivity indices are the mean ± SD of 10 eFAST resamplings, compared with a dummy parameter by one-tailed Welch’s t-test (p < 0.05). Population analyses used N = 1,000 virtual cells, and results are summarized as medians and interquartile ranges.

For single-cell RNA-seq, differential expression was tested per LEC subpopulation using the two-sided Wilcoxon rank-sum test with Benjamini–Hochberg correction. All simulations are deterministic and can be reproduced.

