## Supplementary Information for "Computational modeling of VEGF-A signaling predicts lymphatic junction state"

**This file includes:**

- Supplementary Figures 1–11
- Supplementary Tables 1–3
- Supplementary Note 1: Model formulation (reaction network and ordinary differential equations)
- Supplementary References

**Other supplementary materials for this manuscript (separate files):**

- Supplementary Data 1: Results from every parameter estimation (PSO) run
- Supplementary Data 2: Gamma-distributed initial values for the 1,000 virtual cells

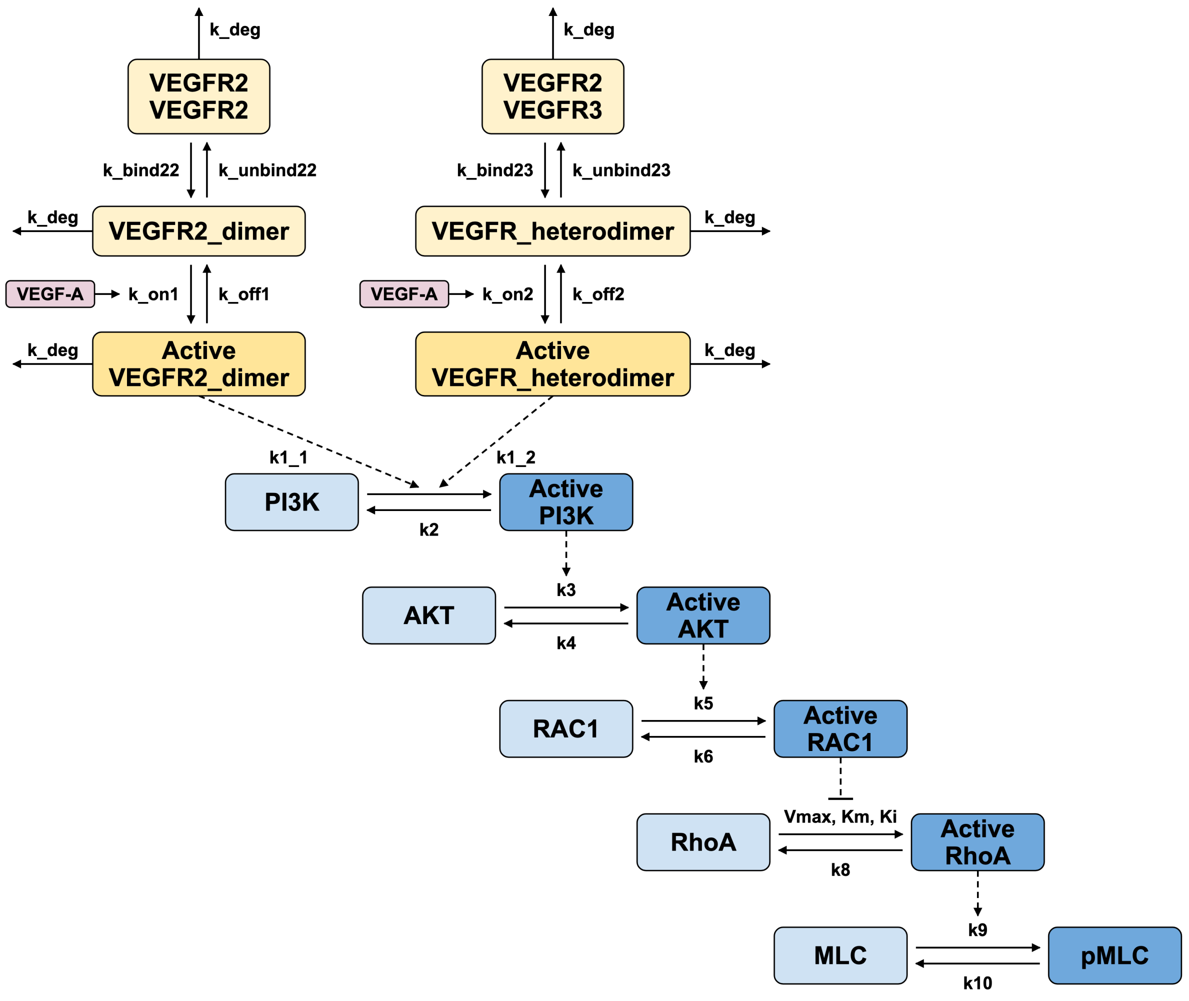


**Supplementary Figure 1. Schematic of the calibrated ODE model.** The signaling network was adapted from Zarkada et al.[1] Pink nodes, ligands; yellow nodes, receptors; blue nodes, downstream signaling species; within each color, darker shading denotes the active form of a species. Solid arrows, molecular conversions; dashed arrows, catalytic or inhibitory regulatory interactions; all parameters are labeled.

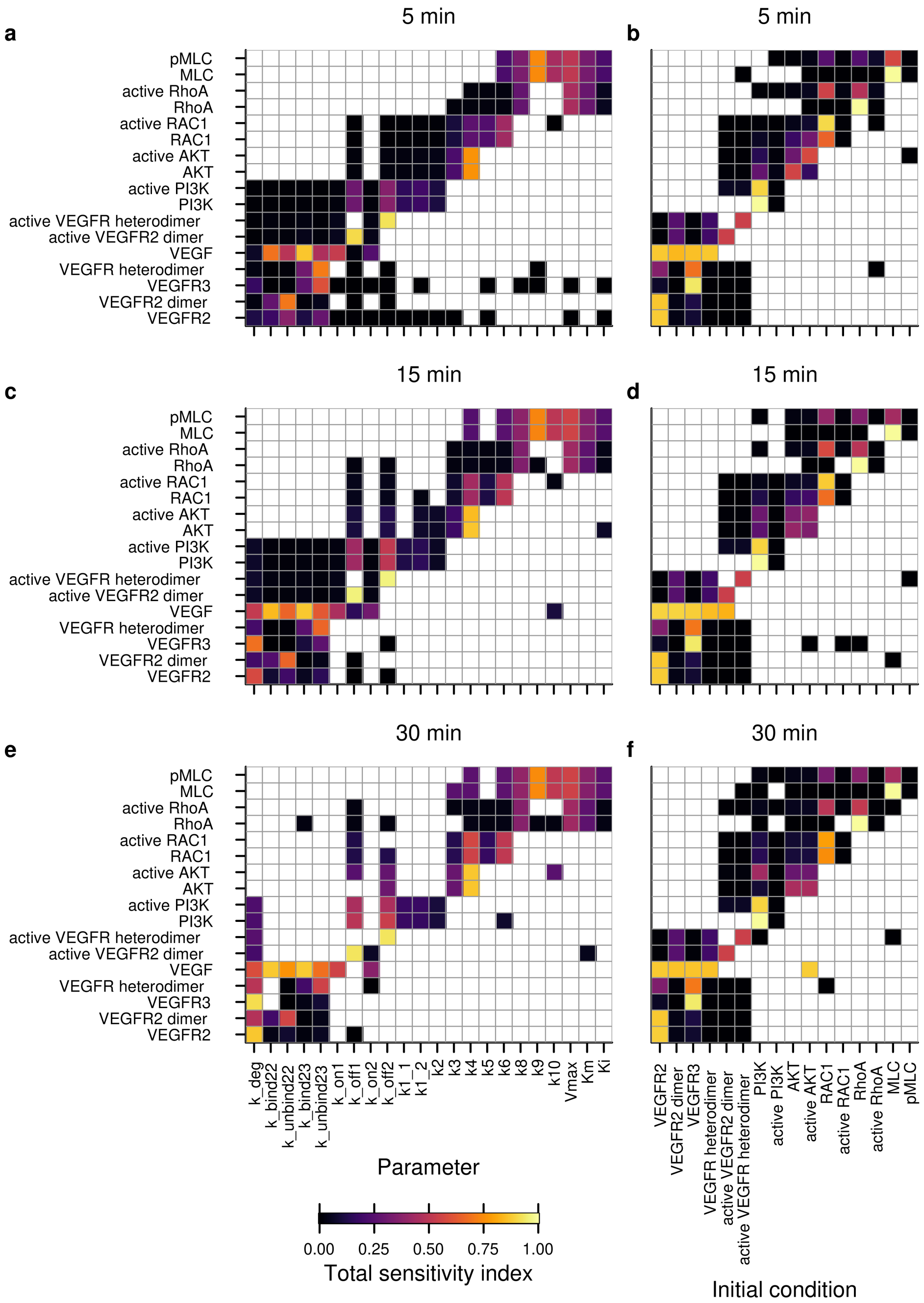


**Supplementary Figure 2. Global sensitivity analysis for model calibration.** Total-order extended Fourier amplitude sensitivity test (eFAST) indices for the 17 model species outputs (rows) with respect to (**a**, **c**, **e**) the 22 kinetic parameters or (**b**, **d**, **f**) the 16 initial conditions (columns), evaluated at $t$ = 5 min (**a**, **b**), 15 min (**c**, **d**), and 30 min (**e**, **f**). Heatmap color indicates the mean of the total-order index (brighter, larger). Blank cells indicate inputs whose index was not statistically distinguishable from that of a dummy parameter.

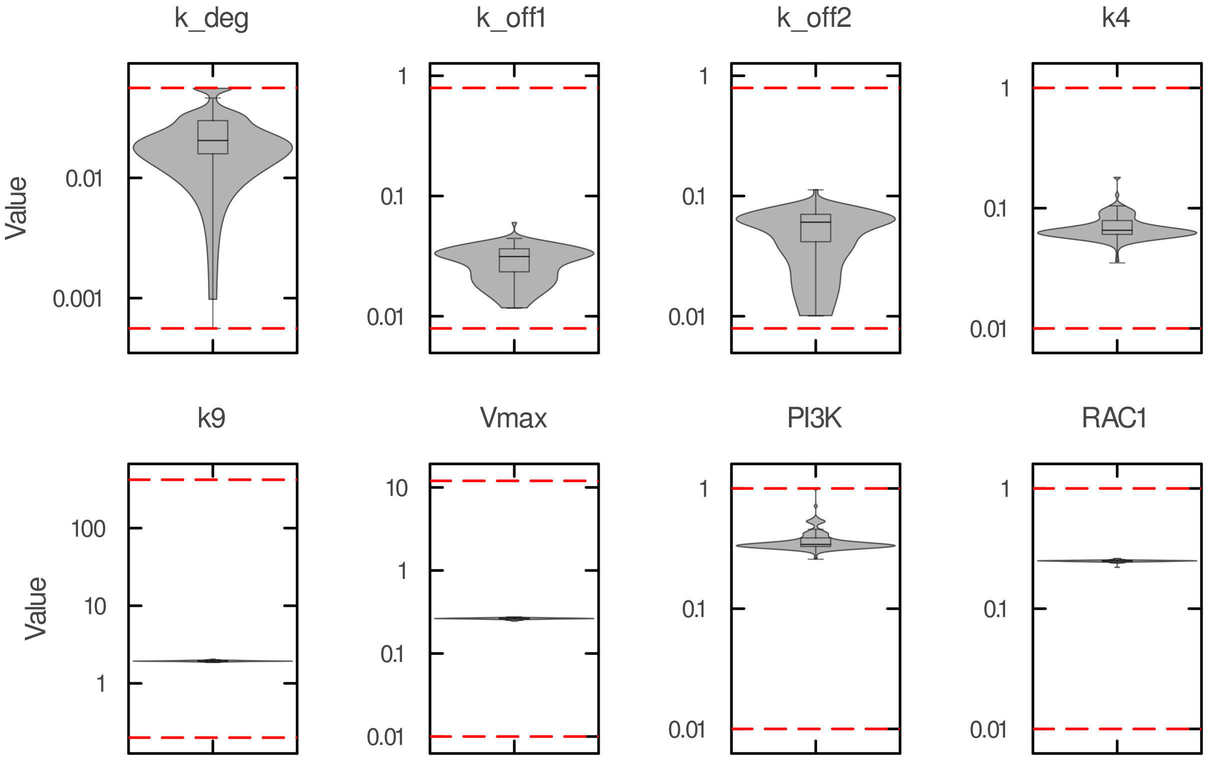


**Supplementary Figure 3. Distribution of parameter estimates across 100 independent particle swarm optimization runs.** Violin plots of each of the eight estimated quantities ($k_{deg}$, $k_{off1}$, $k_{off2}$, $k_4$, $k_9$, $V_{max}$, and the initial values of PI3K and RAC1) across 100 independent particle swarm optimization (PSO) runs, shown on a logarithmic axis. Red dashed lines indicate the lower and upper search bounds for each parameter. Values for every run are provided in Supplementary Data 1.

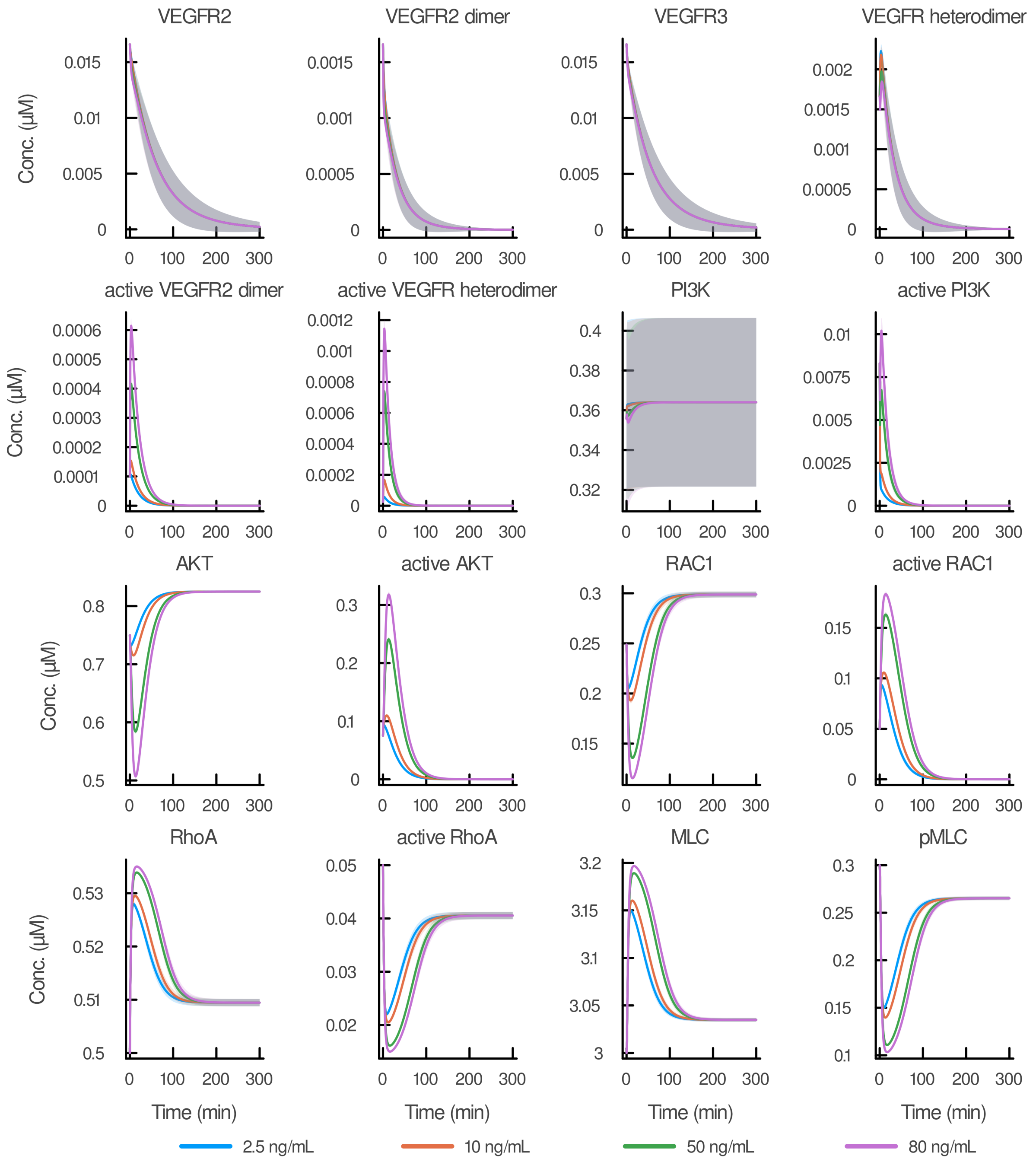


**Supplementary Figure 4. Simulated dynamics of all model species at four VEGF-A doses.** Simulated trajectories of all model species except VEGF-A following stimulation with 2.5, 10, 50, or 80 ng/mL VEGF-A. Lines show the mean over the ensemble of the top 89 of 100 PSO-fitted parameter sets; shaded bands show ± 1 SD across the ensemble; line color indicates dose. PI3K shows a wider spread than other species because its initial value was freely estimated by PSO (Supplementary Fig. 3).

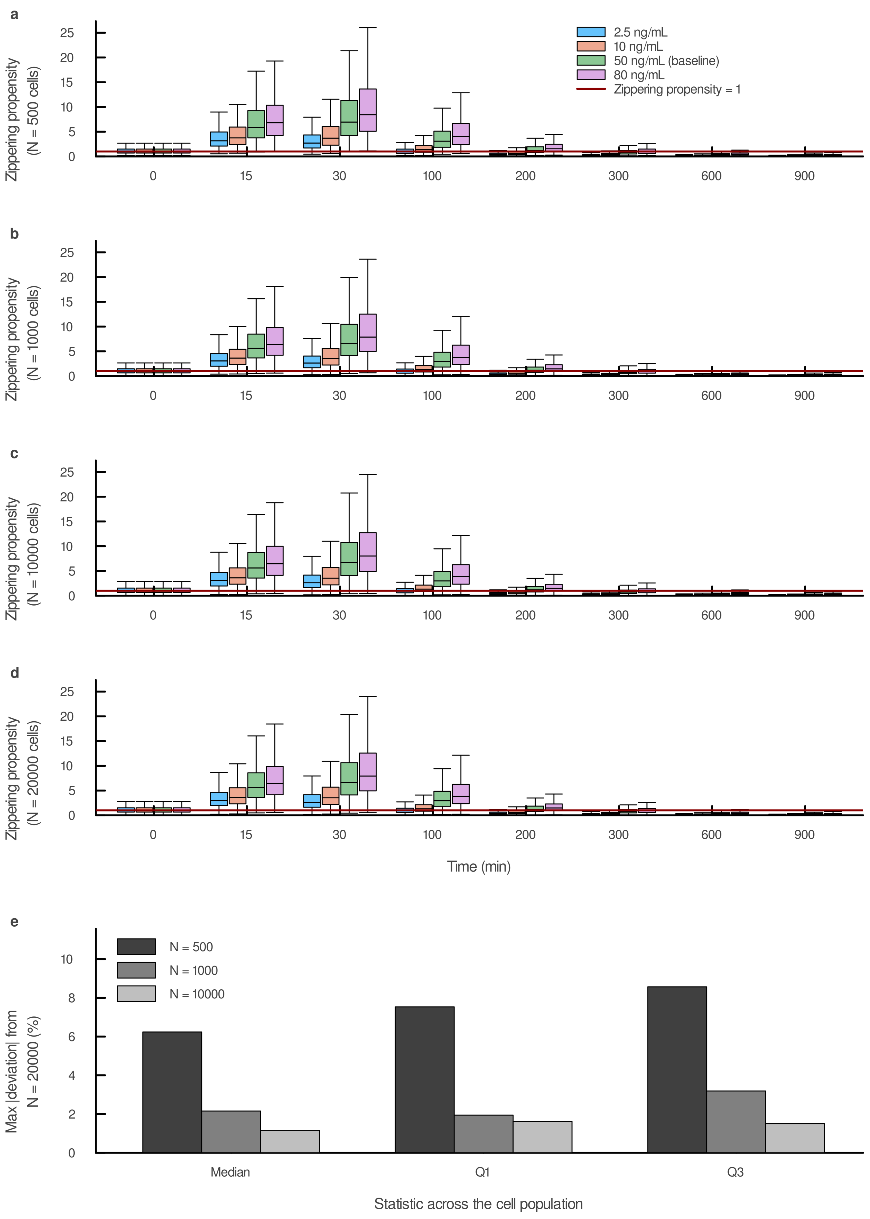


**Supplementary Figure 5. Zippering propensity statistics across population sizes.** (**a-d**) Zippering propensity at eight sampled times (0, 15, 30, 100, 200, 300, 600, and 900 min) at four VEGF-A doses (2.5 ng/mL, blue; 10 ng/mL, orange; 50 ng/mL, green; 80 ng/mL, purple), shown as grouped box plots for heterogeneous populations of $N$ = 500 (**a**), 1,000 (**b**), 10,000 (**c**), and 20,000 (**d**) cells. Boxes span the interquartile range (IQR), the median is marked, and whiskers extend to 1.5× IQR. (**e**) Deviation of the median, first quartile (Q1), and third quartile (Q3) of zippering propensity for populations of $N$ = 500, 1,000, and 10,000 cells from those of the $N$ = 20,000 population, confirming that the $N$ = 1,000 population used throughout this study yields stable estimates.

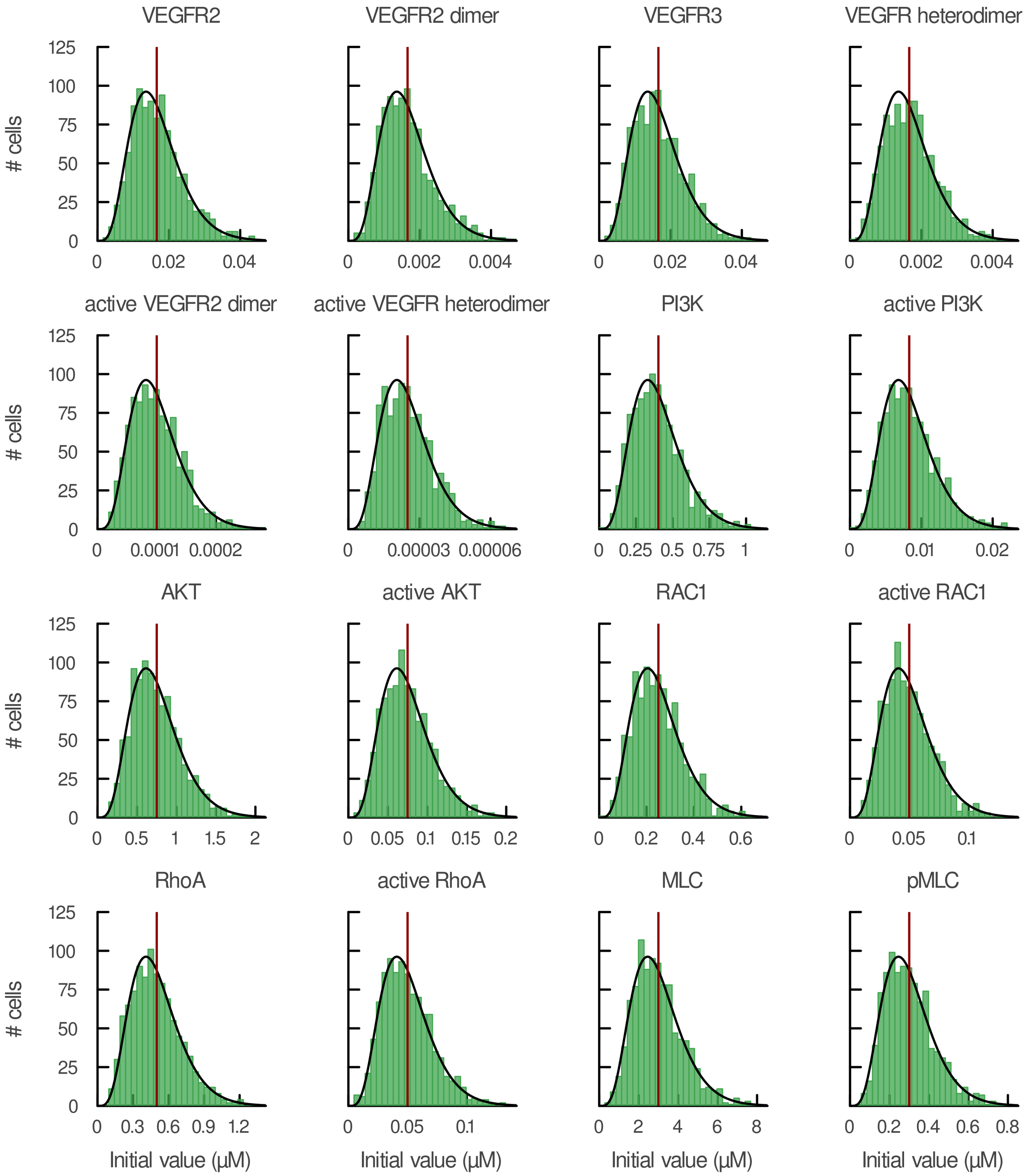


**Supplementary Figure 6. Sampled initial-condition distributions match their target gamma distributions.** Histograms of the sampled initial value of each of the 16 model species across the heterogeneous population ($N$ = 1,000 cells), overlaid with the theoretical gamma probability density function (black line) and the baseline value (dark red line). Each species was sampled from a gamma distribution with shape parameter $a$ = 5.5. Sampled values are provided in Supplementary Data 2.

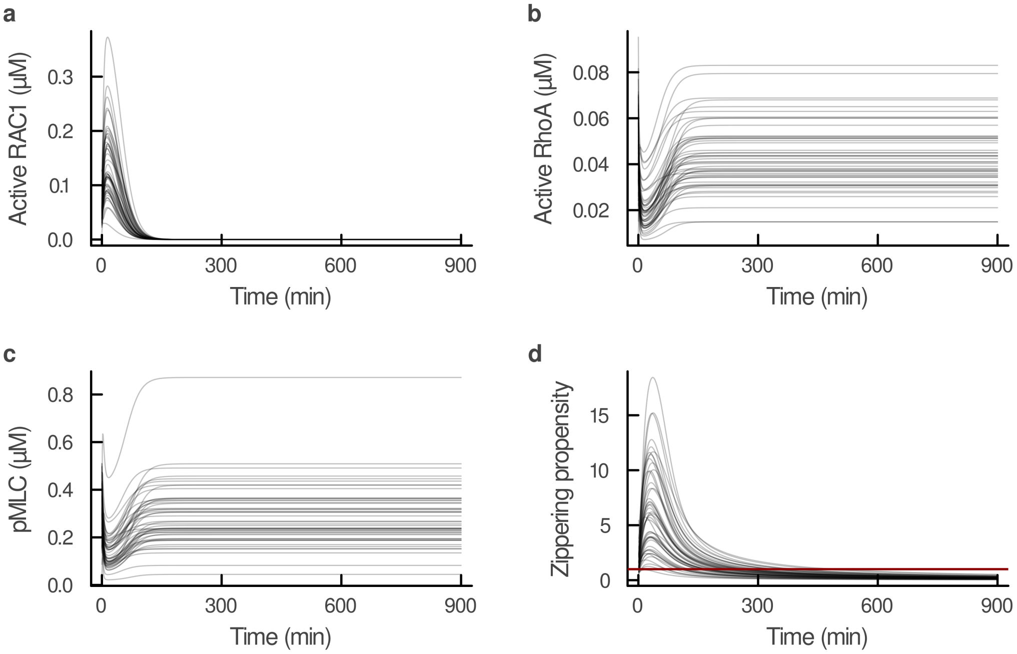


**Supplementary Figure 7. Single-cell trajectories at the calibration dose of VEGF-A.** Simulated dynamics of (**a**) active RAC1, (**b**) active RhoA, (**c**) pMLC, and (**d**) zippering propensity for 50 representative cells from the heterogeneous population ($N$ = 1,000 cells) over 0–900 min following stimulation with 50 ng/mL VEGF-A. The dark red line in **d** marks zippering propensity = 1.

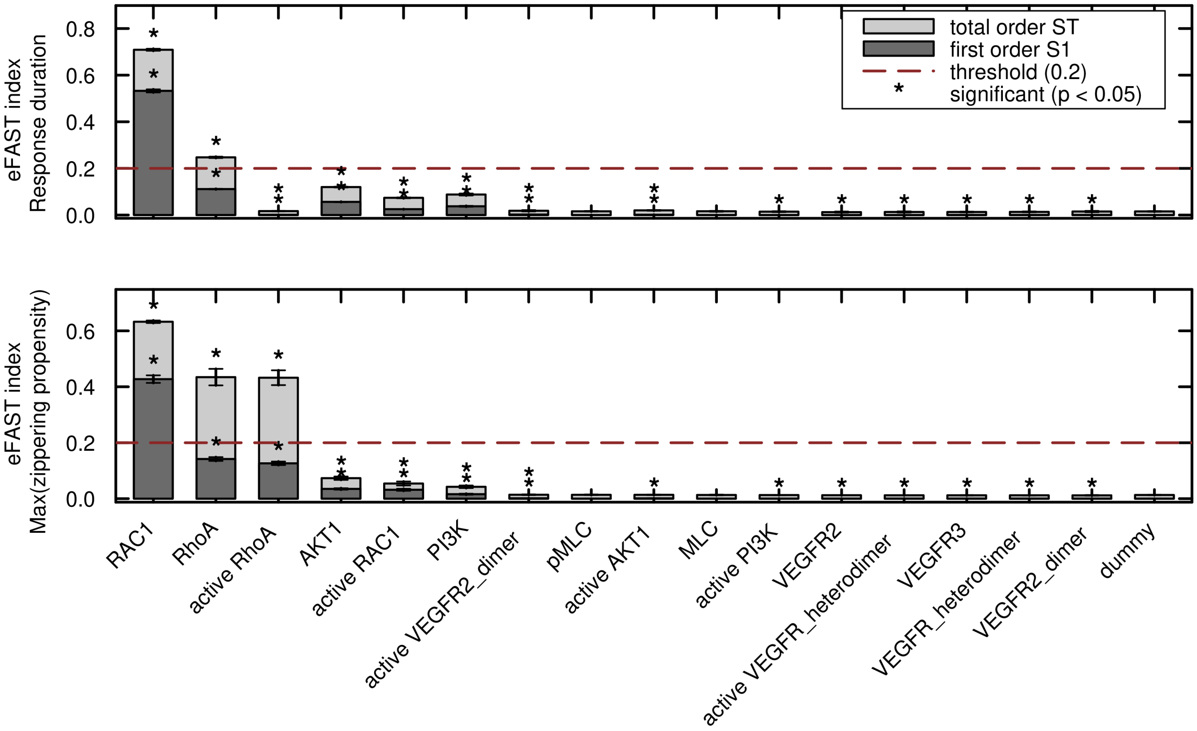


**Supplementary Figure 8. Global sensitivity analysis of initial values in the population-level model.** Total-order (*ST*) and first-order (*S*1) eFAST indices of the model initial values, with max(zippering propensity) and response duration as outputs. Asterisks mark initial values whose indices were significantly different from those of a dummy parameter. Error bars show the SD of 10 eFAST repeats. The influential initial values were RAC1, RhoA, and active RhoA. $N$ = 1,000 cells simulated over 900 min.

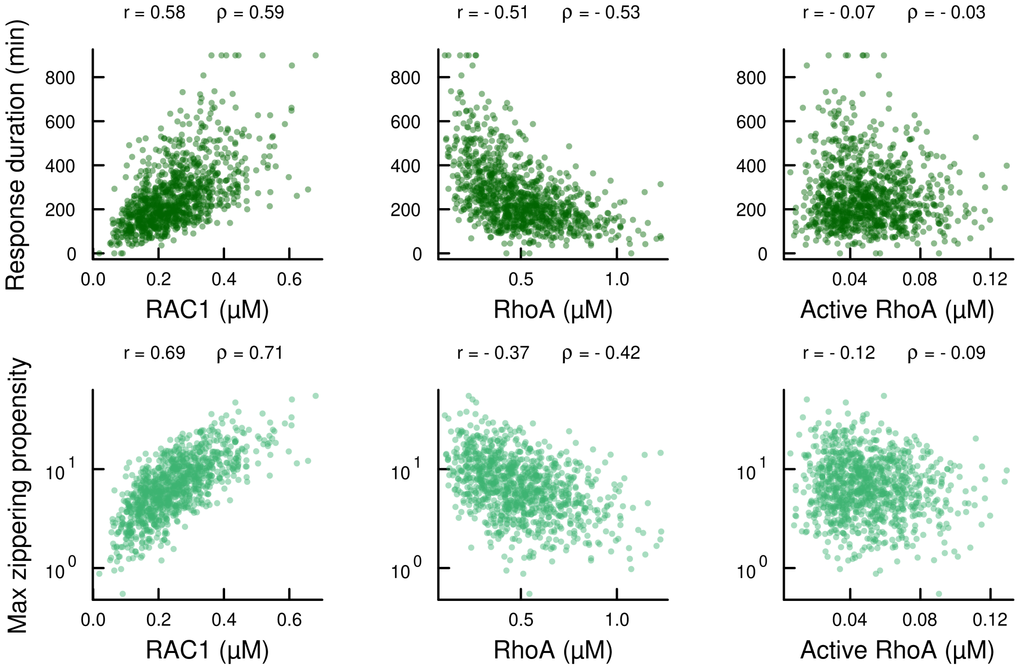


**Supplementary Figure 9. Relationship between influential initial values and model outputs in the population-level model.** Scatter plots of the initial values of RAC1, RhoA and active RhoA, identified as influential in Supplementary Fig. 8, against max(zippering propensity) and response duration. Each point is one simulated cell ($N$ = 1,000 cells); Pearson $r$ and Spearman $\rho$ are shown in each panel.

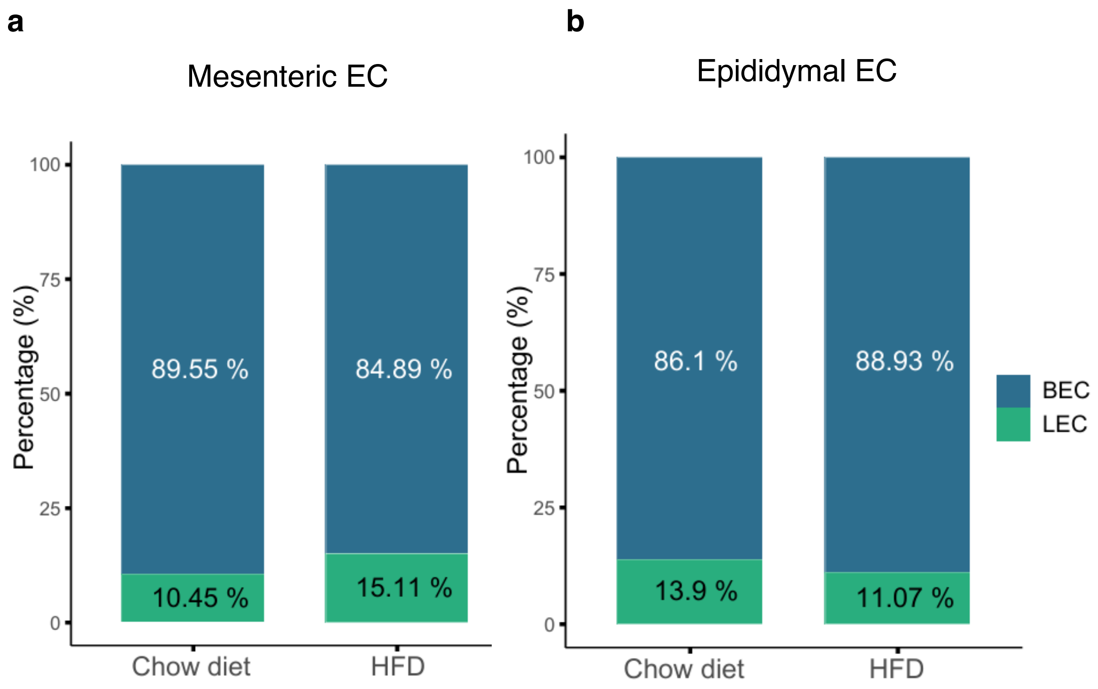


**Supplementary Figure 10. Lymphatic and blood endothelial cell abundance in mesenteric and epididymal tissue under chow and high-fat diet.** Proportions of blood endothelial cells (BECs) and lymphatic endothelial cells (LECs) within the total endothelial compartment of (**a**) mesenteric and (**b**) epididymal tissue from mice fed normal chow or a high-fat diet (HFD), from the single-cell RNA-sequencing dataset of Dunaway et al.[2] In mesenteric tissue, 696 of 6,663 endothelial cells were LECs on chow and 623 of 4,122 on HFD. In epididymal tissue, 1,114 of 8,017 endothelial cells were LECs on chow and 587 of 5,304 on HFD. Counts are cells retained after quality filtering; percentages are of total endothelial cells within each tissue and diet condition.

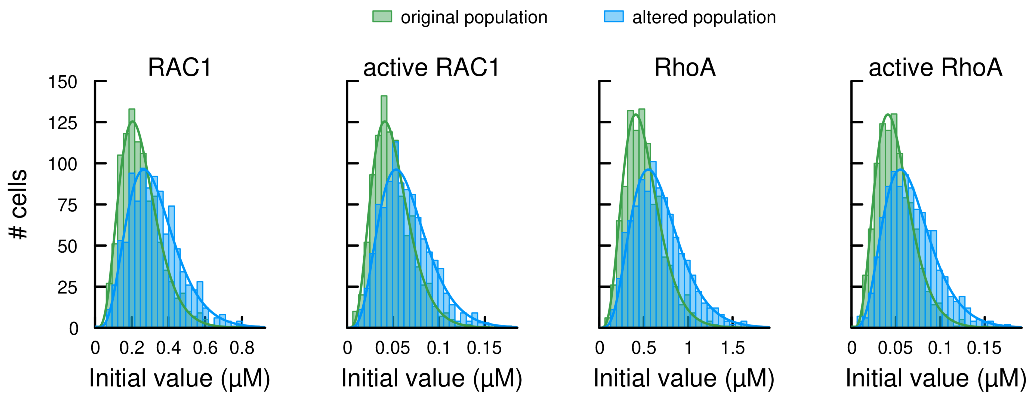


**Supplementary Figure 11. Shifted initial-condition distributions for Model 1 (mesenteric capillary LECs).** Following the perturbation defined in Table 1 of the main text, the initial values distribution of RAC1, active RAC1, RhoA and active RhoA. Each species was resampled from a gamma distribution, Gamma($a$, $b$), with the shape parameter fixed at $a$ = 5.5 for every species and only the mean $m = a \times b$ rescaled (×1.304 for RAC1 and active RAC1; ×1.348 for RhoA and active RhoA). Green, baseline distribution (original populaion); blue, shifted distribution (altered population). $N$ = 1,000 cells sampled.

**Supplementary Table 1. Kinetic parameter values, units, and sources.** Parameters marked "Estimated" were fitted by particle swarm optimization (PSO; Supplementary Data 1); the value shown is from the best-fitting run. $k_{on1}$ and $k_{on2}$ are derived from the fitted $k_{off1}$ and $k_{off2}$ values with a fixed ligand–receptor dissociation constant $K_d = 1.5 \times 10^{-4}$ μM.

| # | Parameter | Biological meaning | Unit | Value | Source |
|---|---|---|---|---|---|
| 1 | $k_{deg}$ | Degradation of all VEGFR species (VEGFR2, VEGFR3, both dimers, both active dimers) | $min^{-1}$ | 0.0313 | Estimated (PSO) |
| 2 | $k_{bind22}$ | VEGFR2 homodimerization | $\mu M^{-1}\,min^{-1}$ | 6 | Fixed[3] |
| 3 | $k_{unbind22}$ | VEGFR2 homodimer dissociation | $min^{-1}$ | 0.6 | Fixed[3] |
| 4 | $k_{bind23}$ | VEGFR2/VEGFR3 heterodimerization | $\mu M^{-1}\,min^{-1}$ | 6 | Fixed[3] |
| 5 | $k_{unbind23}$ | VEGFR2/VEGFR3 heterodimer dissociation | $min^{-1}$ | 0.6 | Fixed[3] |
| 6 | $k_{on1}$ | VEGF-A association to the VEGFR2 homodimer (receptor activation) | $\mu M^{-1}\,min^{-1}$ | 148 | Derived: $k_{off1}/K_d$ |
| 7 | $k_{off1}$ | VEGF-A dissociation from the VEGFR2 homodimer | $min^{-1}$ | 0.0222 | Estimated (PSO) |
| 8 | $k_{on2}$ | VEGF-A association to the VEGFR2/VEGFR3 heterodimer | $\mu M^{-1}\,min^{-1}$ | 256 | Derived: $k_{off2}/K_d$ |
| 9 | $k_{off2}$ | VEGF-A dissociation from the VEGFR2/VEGFR3 heterodimer | $min^{-1}$ | 0.0384 | Estimated (PSO) |
| 10 | $k_{1\_1}$ | PI3K activation by the ligand-bound VEGFR2 homodimer | $\mu M^{-1}\,min^{-1}$ | 30 | Fixed |
| 11 | $k_{1\_2}$ | PI3K activation by the ligand-bound VEGFR2/3 heterodimer | $\mu M^{-1}\,min^{-1}$ | 30 | Fixed |
| 12 | $k_2$ | PI3K deactivation | $min^{-1}$ | 1.8 | Fixed |
| 13 | $k_3$ | AKT phosphorylation by active PI3K | $\mu M^{-1}\,min^{-1}$ | 7.2 | Fixed[4] |
| 14 | $k_4$ | AKT1 dephosphorylation | $min^{-1}$ | 0.0828 | Estimated (PSO) |
| 15 | $k_5$ | RAC1 activation driven by active AKT | $\mu M^{-1}\,min^{-1}$ | 3 | Fixed |
| 16 | $k_6$ | RAC1 inactivation (GTP hydrolysis) | $min^{-1}$ | 0.6 | Fixed |
| 17 | $k_8$ | RhoA inactivation (GTP hydrolysis) | $min^{-1}$ | 0.6 | Fixed |
| 18 | $k_9$ | MLC phosphorylation driven by active RhoA | $\mu M^{-1}\,min^{-1}$ | 1.934 | Estimated (PSO) |
| 19 | $k_{10}$ | pMLC dephosphorylation | $min^{-1}$ | 0.9 | Fixed |
| 20 | $V_{max}$ | Maximal rate of RhoA activation | $\mu M\,min^{-1}$ | 0.264 | Estimated (PSO) |
| 21 | $K_m$ | Michaelis constant of RhoA activation | μM | 5 | Fixed[5] |
| 22 | $K_i$ | Inhibition constant for RAC1-mediated (non-competitive) suppression of RhoA activation | μM | 0.1 | Fixed |

**Supplementary Table 2. Initial values of each model species, units, and sources (baseline).** Values are those of the baseline model. In the heterogeneous population, each species except VEGF-A was sampled from a gamma distribution with shape parameter $a$ = 5.5 and mean equal to the value shown (Supplementary Fig. 6; Supplementary Data 2).

| # | Species | Biological meaning | Unit | Value | Source |
|---|---|---|---|---|---|
| 1 | VEGFR2 | Free VEGFR2 monomer at the LEC surface | μM | 0.0166 | Fixed[3] |
| 2 | VEGFR2_dimer | Unliganded VEGFR2 homodimer | μM | 0.00166 | Fixed[3] |
| 3 | VEGFR3 | Free VEGFR3 monomer | μM | 0.0166 | Fixed[3] |
| 4 | VEGFR_heterodimer | Unliganded VEGFR2/VEGFR3 heterodimer | μM | 0.00166 | Fixed[3] |
| 5 | VEGF | Extracellular VEGF-A stimulus (0.00132 μM = 50 ng/mL, 38 kDa) | μM | 0.00132 | Fixed (stimulus) |
| 6 | Active VEGFR2_dimer | VEGF-bound (active) VEGFR2 homodimer | μM | 0.0001 | Fixed |
| 7 | Active VEGFR_heterodimer | VEGF-bound (active) VEGFR2/3 heterodimer | μM | 0.000025 | Fixed |
| 8 | PI3K | Inactive PI3K | μM | 0.403 | Estimated (PSO) |
| 9 | Active PI3K | Active PI3K | μM | 0.0083 | Fixed[4] |
| 10 | AKT | Unphosphorylated AKT1 | μM | 0.75 | Fixed[4] |
| 11 | Active AKT | Phosphorylated AKT1 | μM | 0.075 | Fixed[4] |
| 12 | RAC1 | GDP-RAC1 (inactive) | μM | 0.25 | Estimated (PSO) |
| 13 | Active RAC1 | GTP-RAC1 (active) | μM | 0.05 | Fixed |
| 14 | RhoA | GDP-RhoA (inactive) | μM | 0.5 | Fixed[5] |
| 15 | Active RhoA | GTP-RhoA (active) | μM | 0.05 | Fixed[5] |
| 16 | MLC | Unphosphorylated myosin light chain | μM | 3 | Fixed[5] |
| 17 | pMLC | Phosphorylated MLC | μM | 0.3 | Fixed[5] |

**Supplementary Table 3. Experimental datasets used for model calibration and validation.** Values are fold change relative to the unstimulated baseline ($t$ = 0) following stimulation with 50 ng/mL VEGF-A. Calibration data were obtained from LECs transfected with control siRNA (si-Ctrl); validation data from LECs with VEGFR3 (si-VEGFR3) or VEGFR2 (si-VEGFR2) knockdown. Active RhoA and active RAC1 were measured at 0 and 30 min only (—, not measured).

| | | | Time (min) | | | |
|---|---|---|---|---|---|---|
| Use | Condition | Model species | 0 | 5 | 15 | 30 |
| Calibration | si-Ctrl | Active AKT | 0.994914 | 2.22549 | 3.43243 | 2.13869 |
| | | Active VEGFR2_dimer | 0.951955 | 4.06946 | 1.90101 | 1.49051 |
| | | pMLC | 0.995696 | 0.466172 | 0.393424 | 0.469195 |
| | | Active RhoA | 0.991707 | — | — | 0.264226 |
| | | Active RAC1 | 0.960484 | — | — | 2.81648 |
| Validation | si-VEGFR3 | Active AKT | 0.99924 | 1.40879 | 2.46540 | 1.50398 |
| | | Active VEGFR2_dimer | 1.00135 | 4.40186 | 1.91228 | 1.49881 |
| | si-VEGFR2 | Active RhoA | 0.98378 | — | — | 0.80394 |
| | | Active RAC1 | 0.93323 | — | — | 0.75763 |

### Supplementary Note 1: Model formulation

#### Reaction network

##### Receptor dimerization.

$$2\,\text{VEGFR2} \rightarrow \text{VEGFR2\_dimer} \qquad (k_{\text{bind22}})$$
$$\text{VEGFR2\_dimer} \rightarrow 2\,\text{VEGFR2} \qquad (k_{\text{unbind22}})$$
$$\text{VEGFR2} + \text{VEGFR3} \rightarrow \text{VEGFR\_heterodimer} \qquad (k_{\text{bind23}})$$
$$\text{VEGFR\_heterodimer} \rightarrow \text{VEGFR2} + \text{VEGFR3} \qquad (k_{\text{unbind23}})$$

##### Ligand–receptor binding.

$$\text{VEGF} + \text{VEGFR2\_dimer} \rightarrow \text{Active VEGFR2\_dimer} \qquad (k_{\text{on1}})$$
$$\text{Active VEGFR2\_dimer} \rightarrow \text{VEGFR2\_dimer} \qquad (k_{\text{off1}})$$
$$\text{VEGF} + \text{VEGFR\_heterodimer} \rightarrow \text{Active VEGFR\_heterodimer} \qquad (k_{\text{on2}})$$
$$\text{Active VEGFR\_heterodimer} \rightarrow \text{VEGFR\_heterodimer} \qquad (k_{\text{off2}})$$

##### Receptor degradation.

$$\text{VEGFR2},\ \text{VEGFR3},\ \text{VEGFR2\_dimer},\ \text{VEGFR\_heterodimer} \rightarrow \emptyset \qquad (k_{\text{deg}})$$
$$\text{Active VEGFR2\_dimer},\ \text{Active VEGFR\_heterodimer} \rightarrow \emptyset \qquad (k_{\text{deg}})$$

##### Downstream signaling.

$$\text{Active VEGFR2\_dimer} + \text{PI3K} \rightarrow \text{Active VEGFR2\_dimer} + \text{Active PI3K} \qquad (k_{1\_1})$$
$$\text{Active VEGFR\_heterodimer} + \text{PI3K} \rightarrow \text{Active VEGFR\_heterodimer} + \text{Active PI3K} \qquad (k_{1\_2})$$
$$\text{Active PI3K} \rightarrow \text{PI3K} \qquad (k_2)$$
$$\text{Active PI3K} + \text{AKT} \rightarrow \text{Active PI3K} + \text{Active AKT} \qquad (k_3)$$
$$\text{Active AKT} \rightarrow \text{AKT} \qquad (k_4)$$
$$\text{Active AKT} + \text{RAC1} \rightarrow \text{Active AKT} + \text{Active RAC1} \qquad (k_5)$$
$$\text{Active RAC1} \rightarrow \text{RAC1} \qquad (k_6)$$
$$\text{RhoA} \rightarrow \text{Active RhoA} \qquad (v_7)$$
$$\text{Active RhoA} \rightarrow \text{RhoA} \qquad (k_8)$$
$$\text{Active RhoA} + \text{MLC} \rightarrow \text{Active RhoA} + \text{pMLC} \qquad (k_9)$$
$$\text{pMLC} \rightarrow \text{MLC} \qquad (k_{10})$$

where RhoA activation follows Michaelis–Menten kinetics with non-competitive inhibition by active RAC1,

$$v_7 = \frac{V_{\text{max}}\,[\text{RhoA}]}{(K_m + [\text{RhoA}])(1 + [\text{Active RAC1}]/K_i)}.$$

#### Model ODEs

$$\frac{d[\text{VEGFR2}]}{dt} = -k_{\text{deg}}[\text{VEGFR2}] - k_{\text{bind22}}[\text{VEGFR2}]^2 + 2k_{\text{unbind22}}[\text{VEGFR2\_dimer}] - k_{\text{bind23}}[\text{VEGFR2}][\text{VEGFR3}] + k_{\text{unbind23}}[\text{VEGFR\_heterodimer}]$$

$$\frac{d[\text{VEGFR2\_dimer}]}{dt} = -k_{\text{deg}}[\text{VEGFR2\_dimer}] + \tfrac{1}{2}k_{\text{bind22}}[\text{VEGFR2}]^2 - k_{\text{unbind22}}[\text{VEGFR2\_dimer}] - k_{\text{on1}}[\text{VEGF}][\text{VEGFR2\_dimer}] + k_{\text{off1}}[\text{Active VEGFR2\_dimer}]$$

$$\frac{d[\text{VEGFR3}]}{dt} = -k_{\text{deg}}[\text{VEGFR3}] - k_{\text{bind23}}[\text{VEGFR2}][\text{VEGFR3}] + k_{\text{unbind23}}[\text{VEGFR\_heterodimer}]$$

$$\frac{d[\text{VEGFR\_heterodimer}]}{dt} = -k_{\text{deg}}[\text{VEGFR\_heterodimer}] + k_{\text{bind23}}[\text{VEGFR2}][\text{VEGFR3}] - k_{\text{unbind23}}[\text{VEGFR\_heterodimer}] - k_{\text{on2}}[\text{VEGF}][\text{VEGFR\_heterodimer}] + k_{\text{off2}}[\text{Active VEGFR\_heterodimer}]$$

$$\frac{d[\text{VEGF}]}{dt} = -k_{\text{on1}}[\text{VEGF}][\text{VEGFR2\_dimer}] - k_{\text{on2}}[\text{VEGF}][\text{VEGFR\_heterodimer}]$$

$$\frac{d[\text{Active VEGFR2\_dimer}]}{dt} = -k_{\text{deg}}[\text{Active VEGFR2\_dimer}] + k_{\text{on1}}[\text{VEGF}][\text{VEGFR2\_dimer}] - k_{\text{off1}}[\text{Active VEGFR2\_dimer}]$$

$$\frac{d[\text{Active VEGFR\_heterodimer}]}{dt} = -k_{\text{deg}}[\text{Active VEGFR\_heterodimer}] + k_{\text{on2}}[\text{VEGF}][\text{VEGFR\_heterodimer}] - k_{\text{off2}}[\text{Active VEGFR\_heterodimer}]$$

$$\frac{d[\text{Active PI3K}]}{dt} = k_{1\_1}[\text{Active VEGFR2\_dimer}][\text{PI3K}] + k_{1\_2}[\text{Active VEGFR\_heterodimer}][\text{PI3K}] - k_2[\text{Active PI3K}]$$

$$\frac{d[\text{PI3K}]}{dt} = -k_{1\_1}[\text{Active VEGFR2\_dimer}][\text{PI3K}] - k_{1\_2}[\text{Active VEGFR\_heterodimer}][\text{PI3K}] + k_2[\text{Active PI3K}]$$

$$\frac{d[\text{Active AKT}]}{dt} = k_3[\text{Active PI3K}][\text{AKT}] - k_4[\text{Active AKT}]$$

$$\frac{d[\text{AKT}]}{dt} = -k_3[\text{Active PI3K}][\text{AKT}] + k_4[\text{Active AKT}]$$

$$\frac{d[\text{Active RAC1}]}{dt} = k_5[\text{Active AKT}][\text{RAC1}] - k_6[\text{Active RAC1}]$$

$$\frac{d[\text{RAC1}]}{dt} = -k_5[\text{Active AKT}][\text{RAC1}] + k_6[\text{Active RAC1}]$$

$$\frac{d[\text{Active RhoA}]}{dt} = v_7 - k_8[\text{Active RhoA}]$$
$$\frac{d[\text{RhoA}]}{dt} = -v_7 + k_8[\text{Active RhoA}]$$
$$\frac{d[\text{pMLC}]}{dt} = k_9[\text{Active RhoA}][\text{MLC}] - k_{10}[\text{pMLC}]$$
$$\frac{d[\text{MLC}]}{dt} = -k_9[\text{Active RhoA}][\text{MLC}] + k_{10}[\text{pMLC}]$$
